# Host cell potassium ion channels KCNJ2 (K_IR_2.1), KCNJ13 (K_IR_7.1) and KCNMA1 (BK_Ca_) mediate escape of Bunyamwera virus from late endosomal compartments

**DOI:** 10.64898/2026.01.12.698685

**Authors:** Hayley M. Pearson, Samantha Hover, Eleanor J. A. A. Todd, Kyriakoulla Panayi, Martin Stacey, Jamel Mankouri, Jonathan D. Lippiat, Eric W. Hewitt, John N. Barr

## Abstract

The *Orthobunyavirus* genus within the *Peribunyaviridae* family of enveloped arthropod-borne negative-sense RNA viruses includes species associated with serious or fatal disease in both animals and humans such as Schmallenberg, La Crosse and Oropouche viruses. Orthobunyaviruses (OBVs) are internalised into cells by endocytosis and release their genomes following fusion with late endosomal (LE) membranes, triggered by low pH of the luminal milieu. There is mounting evidence to suggest OBV endosomal escape is also influenced by potassium ions (K^+^), which increase in concentration as endosomes mature. Endosomal K^+^ flux is controlled by cellular K^+^ channels, and we previously showed that K^+^ channel blockade using broad spectrum pharmacological inhibition abrogates OBV infection, with virions trapped within the endosomes. However, the K^+^ channels regulating this process are unknown. Herein, to identify the K^+^ channels involved, we studied Bunyamwera virus (BUNV), the prototypical OBV, and screened a siRNA library targeting 342 human ion channels. We identified 19 K^+^ channels whose knockdown inhibited BUNV gene expression by over 50%, with at least seven channels affecting BUNV at the entry stage, suggesting they exert a combinatorial influence. Of these seven channels, we used both pharmacological inhibition and genetic channel manipulation to show that OBV escape from CD63- and Rab7-positive LEs is controlled by KCNJ2 (K_IR_2.1), KCNJ13 (K_IR_7.1) and KCNMA1 (BK_Ca_). These studies add to the understanding of host factors influencing OBV entry, regulation of endosomal ion flux, and could aid in the design or repurposing of therapeutics for the treatment of OBV-associated disease.

## Introduction

The *Orthobunyavirus* genus within the *Peribunyaviridae* family and *Bunyaviricetes* class comprises enveloped arthropod-borne segmented negative-sense RNA viruses, many of which are associated with serious or fatal disease in both animals and humans. Bunyamwera virus (BUNV) is the prototypical orthobunyavirus (OBV) and while it is mostly associated with a mild febrile illness in humans, the BUNV reassortant Ngari virus has been associated with fatal human disease, including haemorrhagic fever (Heitmann et al., 2021). Other serious pathogens within the OBV group includes Oropouche virus (OROV), capable of vertical mother to fetus transmission and associated with stillbirth and birth defects including microcephaly, and La Crosse virus, responsible for widespread outbreaks of encephalitis, with rare but serious neurological complications. In addition, several OBVs inflict a significant impact on animal health, including Aino virus, Akabane virus and Schmallenberg virus (SBV), associated with severe reproductive losses and congenital malformations.

Orthobunyaviruses encode three genomic RNA segments termed small (S), medium (M) and large (L). The S segment encodes the nucleocapsid protein (NP) and the non-structural NSs protein, the M segment encodes a polyprotein precursor that is cleaved to yield glycoproteins Gn and Gc along with non-structural protein NSm, and the L segment encodes the large protein (L), which forms the RNA-dependent RNA polymerase (RdRp). These RNA segments are encapsidated by NP and complexed with the RdRp to form pseudo-circular helical ribonucleoproteins (RNPs). During assembly, these RNPs are packaged within a host-derived envelope through which protrude tripodal glycoprotein spikes, comprising trimers of Gn/Gc heterodimers.

Key stages of the mammalian cell infection cycle are conserved between OBVs, including clathrin-mediated internalisation and trafficking through the endocytic pathway, with virus fusion triggered in late endosomal compartments in response to their precise biochemical composition. Critical cues include a low pH of around 5.0-6.0 (Windhaber et al., 2022), and a high concentration of potassium (K^+^) ions, which together induce the fusion of the endosomal and viral membranes, primarily mediated by Gc, a type II fusion protein (Hellert et al., 2023), with subsequent release of the infecting RNPs into the cytosol.

We have previously demonstrated a critical role of K^+^ during OBV entry and fusion at the late endosome (LE). We showed broad spectrum pharmacological inhibition of K^+^ channels disrupted endosomal K^+^ accumulation and, in turn, prevented the release of BUNV from the LE compartment (Hover et al., 2016, 2018). Instead, unfused viruses were trafficked deeper within the endocytic pathway to lysosomes, resulting in the degradation of the virion and a lack of productive infection. Furthermore, incubation of BUNV prior to incubation with cells, at low pH and high K^+^ concentration ([K^+^]) was sufficient to expedite infection due to a dramatic rearrangement of the glycoprotein trimer, shown through cryo-electron tomography (Hover et al., 2023), revealing that acidification is not the sole trigger required for BUNV entry. BUNV is not the only OBV that depends on K^+^ for entry, with recent reports showing infection with SBV (Hover et al., 2016), Germiston virus (Windhaber et al., 2022), LACV and Keystone virus (Sandler et al., 2020) also depend on maintenance of a K^+^ gradient. Furthermore, we and others have shown viruses from other families within the *Bunyaviricetes* class also depend on elevated K^+^ concentrations for virus multiplication including the *Arenaviridae* family (Shaw et al., 2024), the *Nairoviridae* family (Charlton et al., 2019; Punch et al., 2018) and the *Phenuiviridae* family (Sandler et al., 2020). The finding that different families are susceptible to pharmacological inhibition of K^+^ channels suggests a shared mechanism, which could be targeted therapeutically by pan-bunyaviral treatments.

Here, we used BUNV to determine the molecular identity of cellular K^+^ channels that support OBV infection. By siRNA-mediated knockdown of 342 host cell ion channel genes, we showed that control of K^+^ flux relevant to OBV infection involves multiple channels, with knockdown of at least 19 different K^+^ channels significantly inhibiting BUNV gene expression by over 50%. Of these, at least seven K^+^ channels from three different channel families were required for virus entry, and we provide evidence using both pharmacological inhibition and genetic channel manipulation that three of these, namely KCNJ2 (K_IR_2.1), KCNJ13 (K_IR_7.1) and KCNMA1 (BK_Ca_) influence BUNV escape from CD63- and Rab7-positive LE. These studies add to our understanding of host-OBV interactions, the composition of endosomal channelome and furthermore, could ultimately aid in the design or repurposing of therapeutics for the treatment of OBV-associated disease.

## Results

### Identification of human ion channels required during BUNV infection

Our previous work has emphasised the importance of cellular K^+^ channels during BUNV infection, with broad spectrum K^+^ channel blockade preventing virion uncoating within the endocytic network (Hover et al., 2016, 2018). To confirm the identity of the specific K^+^ channel(s) required, we used a gene expression knockdown screen in BUNV-infected human origin A549 cells, targeting 342 host cell ion channel genes with 3 unique siRNAs per gene. To facilitate this analysis, we generated a recombinant BUNV that expressed eGFP (rBUNV-eGFP), allowing live cell analysis of eGFP expression as a measure of viral protein expression.

This recombinant virus was designed such that its S segment mRNA encoded tandem eGFP and BUNV NP ORFs, separated by a porcine teschovirus-1 2A peptide linker (P2A), allowing expression of eGFP and BUNV NP as independent entities. Corresponding plasmid pBUNV-S-eGFP (Fig 1A) was incorporated within a previously described BUNV rescue system(Bridgen & Elliott, 1996; Lowen et al., 2004), with successful rescue of rBUNV-eGFP confirmed by western blotting (Fig 1B, C) and live cell imaging (Fig 1E). Plaque and focus-forming assays were used for growth quantification, with titres and lesion size being comparable between both viruses (Fig 1D, E).

**Figure 1.**
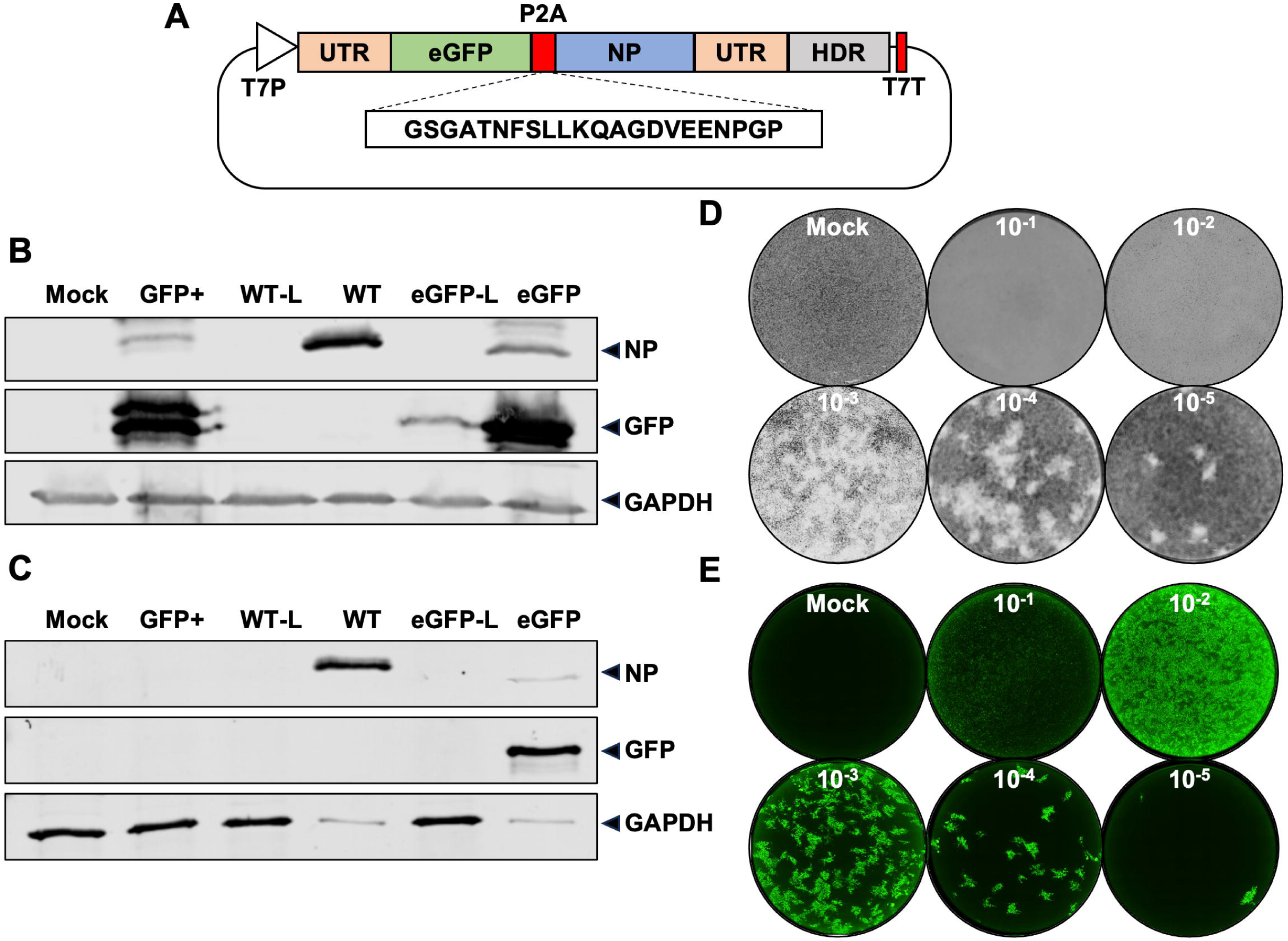
Generation of recombinant BUNV expressing enhanced green fluorescent protein. A. Schematic representation of pBUNV-S-eGFP, encoding a modified BUNV S segment designed to express eGFP and NP ORFs separated by a porcine teschovirus-1 2A peptide linker (P2A). Also shown are the T7 RNA polymerase promoter and terminator signals (T7P/T7T), the untranslated regions (UTR), and the hepatitis delta virus ribozyme (HDR). B. BSR-T7 cells were transfected with either WT S, M and L BUNV rescue plasmids, or with S-eGFP, M and L to rescue rBUNV-eGFP. Controls lacking the L plasmid were included (-L), as well as a transfection control plasmid (GFP+). At 5 days post-transfection, cells were lysed and analysed by western blot for both NP and eGFP expression. C. Supernatants from the transfected BSR-T7 cells were transferred to naïve BHK-21 cells and incubated for 2 days. Cells were lysed and analysed by western blot for NP and eGFP expression. D-E. Supernatant was harvested from the BHK-21 cells and titrated via a plaque assay. rBUNV-WT plaques were visualised by crystal violet staining (D) and rBUNV-eGFP fluorescent plaques were visualised by IncuCyte S3 (E).

The ability of the three unique siRNAs per gene to inhibit rBUNV-eGFP growth was measured by assessing the total integrated intensity of eGFP (TIIE) signal at 24 hours post infection (hpi) using a reverse transfection protocol (Fig. 2A). TIIE was assessed in quadruplicate, with a mean score calculated for each of the three siRNAs. To display the relative effectiveness of each gene knockdown on rBUNV-eGFP growth, the channels knocked down were ranked according to the median score of the three corresponding siRNAs. This revealed knockdown of 46 of the 342 gene targets resulted in greater than a 50% reduction in rBUNV-eGFP signal compared to the transfection reagent-only control (Supp Fig 1). Of these 46 channels, 19 (41%) were K^+^ channels (Fig 2B) consistent with our previous studies showing pharmacological disruption of K^+^ channel activity abrogated BUNV infection.

**Figure 2.**
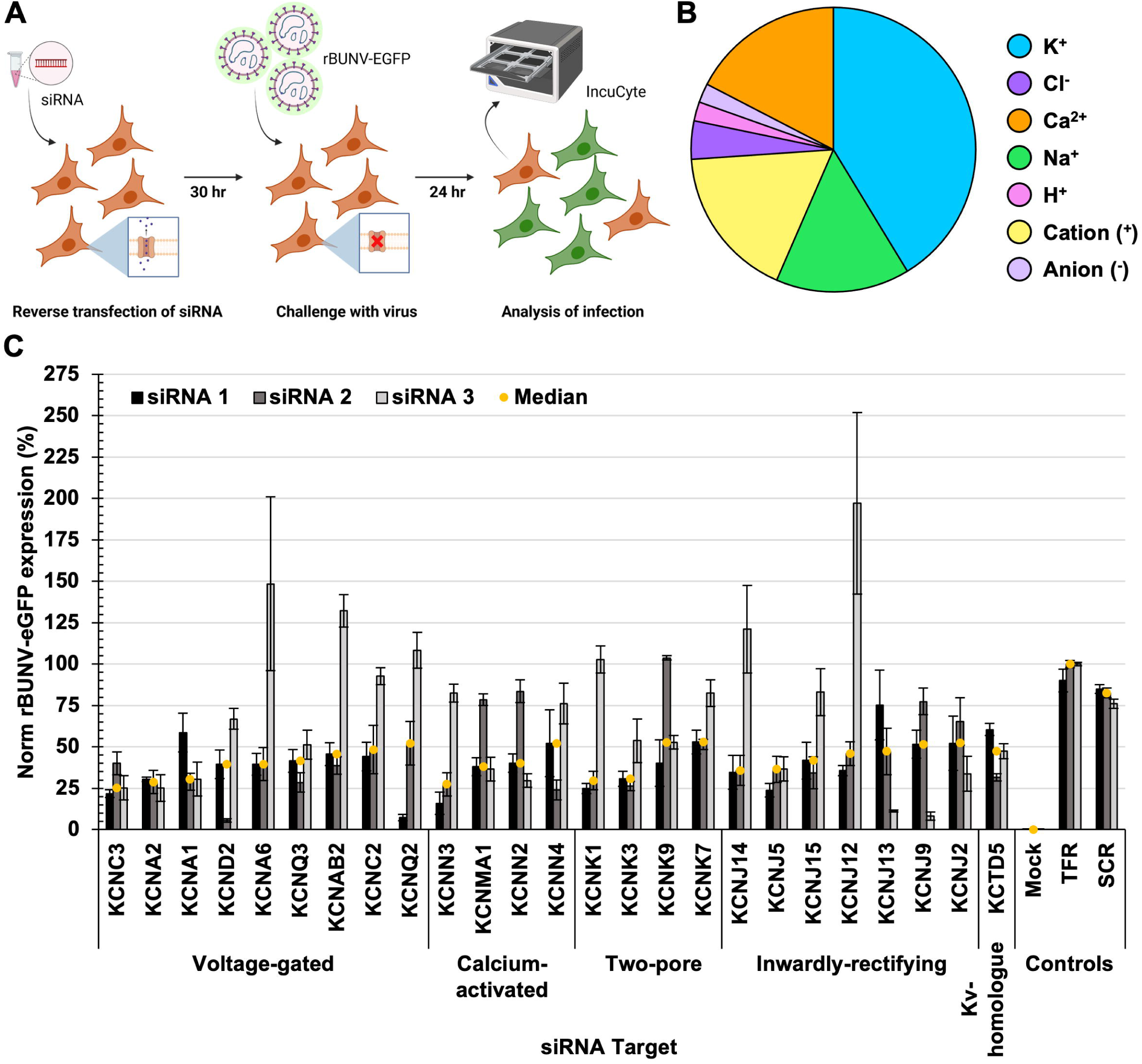
Identification of human ion channels required during BUNV infection using siRNA screening. A. Schematic representation of siRNA screening method. A549 cells were reverse transfected with siRNA targeting human ion channel genes for 30 hrs, then challenged with rBUNV-eGFP. Infection was assessed after 24 hrs by quantification of total integrated intensity of green fluorescence by IncuCyte S3. B. Pie chart of ion channels which inhibited rBUNV-eGFP expression by >50% separated by ion selectivity. C. Top 25 K^+^ channel siRNA target hits based on total integrated intensity of rBUNV-eGFP signal across three unique siRNAs (black, dark grey, light grey bars), grouped by channel family. Each bar represents the mean of four experimental repeats, with the standard error shown. The median value for each target is shown by a yellow circle.

The K^+^ channels identified as affecting rBUNV-eGFP activities were from multiple channel families, with the top-25 ranked K^+^ channel genes (Fig 2C) including 9 voltage-gated K^+^ channels (K_v_), 4 calcium (Ca^2+^)-activated K^+^ channels (K_Ca_), 4 two-pore K^+^ channels (K_2P_), 7 inwardly rectifying K^+^ channels (K_ir_), and 1 K_V_ homologue. The finding that siRNA-mediated knockdown of numerous channels affected BUNV infection suggests several channels may influence K^+^ influx in relation to BUNV multiplication through combinatorial affects.

We next tested whether each of the top-25 ranked ion channel gene hits were expressed in A549 cells. Total RNA was isolated from A549 cells and then amplified using RT-qPCR with specific primer pairs against the K^+^ subunit genes alongside GAPDH for reference. We detected expression of all 25 K^+^ channel targets (Supp Fig 2A) indicating that the siRNA effects observed on the virus were likely due to the knockdown of the target genes.

### Inhibition of select K_Ca_ channels blocks BUNV endosomal escape

To better understand the role of the top-ranked K^+^ channels during BUNV infection, we performed pharmacological inhibition at defined times throughout the infection cycle. Small molecule inhibitors against the 4 Ca^2+^-activated K+ channel hits (Fig 3A) were tested for their impact on infection. A549 cells were pre-treated with each inhibitor or a solvent control prior to infection with rBUNV-eGFP, with infection assessed through the detection of eGFP fluorescence at 24 hpi. Alongside the virus assays, MTS assays were also performed to establish non-cytotoxic concentrations of each compound (>80% cell viability compared to solvent controls).

**Figure 3.**
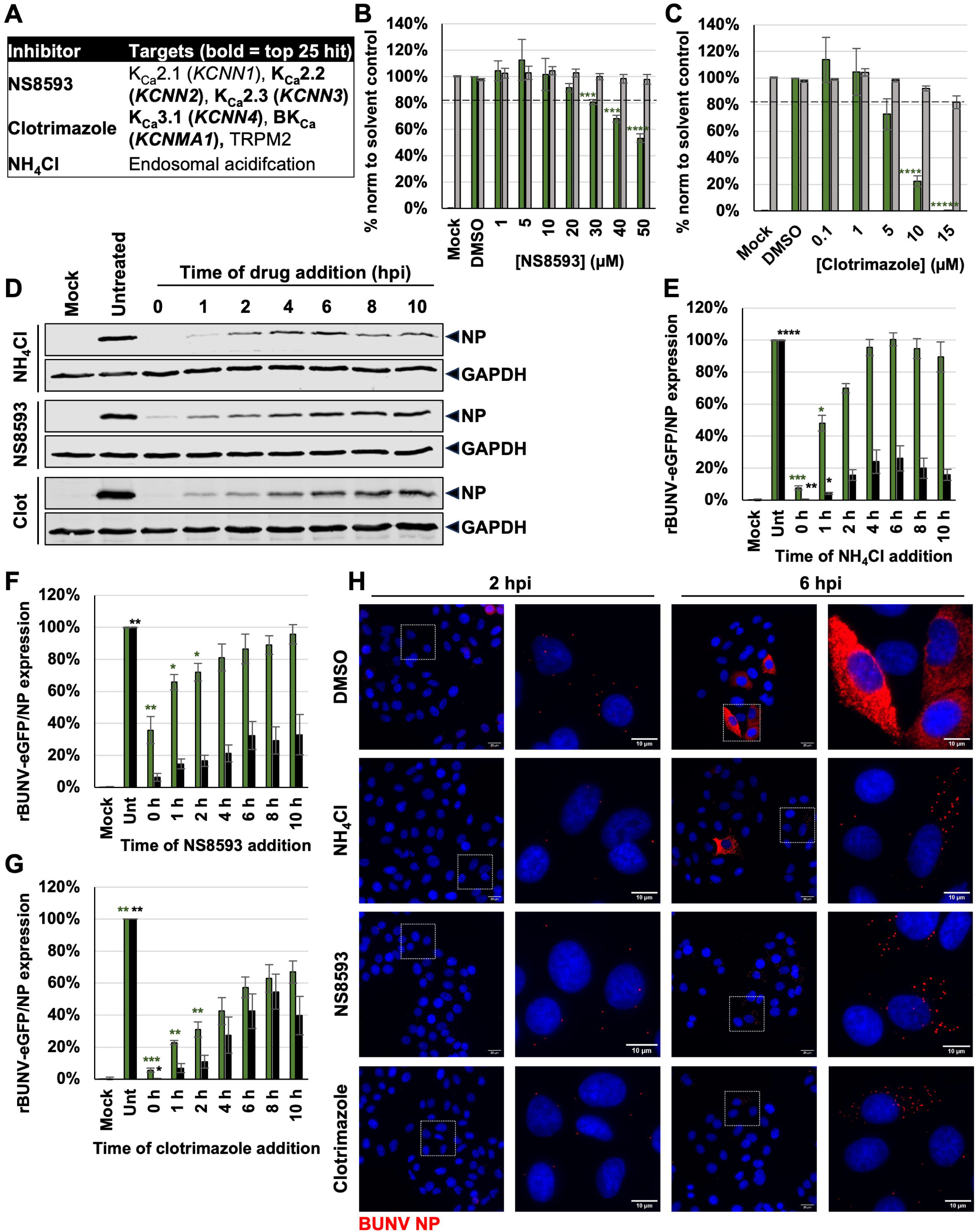
Inhibition of select K_Ca_ channels block BUNV endosomal escape. A. Table to show targets of small molecular inhibitors targeting K_Ca_ channels. Targets in bold were hits in the siRNA screen. Channel protein names are shown, with gene names in brackets. B-C. A549 cells were pre-treated with small molecule inhibitors in A. After 45 mins of treatment, cells were infected with rBUNV-eGFP at MOI 0.1 for 24 hours. The number of rBUNV-eGFP expressing cells was quantified using IncuCyte S3 live imaging and normalised to the solvent control (green bars). Averages of three biological repeats are shown, each performed in duplicate. Drug cytotoxicity was measured immediately afterward via MTS assay and normalised to solvent controls (grey bars), The dashed line represents the toxicity threshold. Three biological repeats were performed, each in triplicate. D-G. A549 cells were infected with either rBUNV-WT or rBUNV-eGFP at MOI 0.5 for 24 hours. At the indicated time points post-infection, infected cells were treated with small molecule inhibitors of K^+^ channels. Cells in the 0 hpi condition were pre-treated for 45 mins prior to infection. D. Representative western blot analysis of BUNV NP and GAPDH expression. E-G. Densitometry analysis of rBUNV-WT NP expression (black bars) alongside number of rBUNV-eGFP expressing cells quantified by IncuCyte S3 (green bars), normalised to untreated (solvent) controls. Data is averaged from three biological repeats. One way ANOVA tests were performed to assess statistical significance. P< 0.05*, P<0.01**, P<0.001***, P<0.0001****. H. A549 cells were pre-treated with the indicated K^+^ channel small molecule inhibitors for 45 mins. Cells were incubated on ice and rBUNV-WT was added at MOI 5 for 1 hour to promote virus binding but not internalisation. Non-internalised virus was removed, and infection was allowed to progress in the presence of the inhibitor at 37°C. Cells were fixed at 2 or 6 hpi and immuno-stained for BUNV NP (red) and DAPI (blue). Solvent control (DMSO) was included, as was internalisation control (NH_4_Cl). Images were taken on Olympus IX83 widefield deconvolution microscope at 60x magnification. White boxes indicate zoomed field of view. Scale bars are included (20 µM wide, 10 µM zoom).

Treatment with 50 µM compound NS8593, which inhibits small conductance Ca^2+^-activated K^+^ channels *KCNN2* (SK2/K_Ca_2.2) and *KCNN3* (SK3/K_Ca_2.3) (Strøbæk et al., 2006), significantly decreased the number of rBUNV-eGFP infected cells to 53% of the solvent control (Fig 3B). Similarly, treatment with 15 µM clotrimazole, which inhibits both *KCNN4* (K_Ca_3.1) and large-conductance Ca^2+^-activated K^+^ channel *KCNMA1* (BK_Ca_/K_Ca_1.1) (Ishii et al., 1997; Rittenhouse et al., 1997; Wu et al., 1999) resulted in complete abrogation of rBUNV-eGFP expression in treated cells (Fig 3C).

Next, we used NS8593 and clotrimazole in time-of-addition experiments to determine at which stage in BUNV infection cycle the K_Ca_ channels are required. For this, rBUNV-WT or rBUNV-eGFP were used to infect A549 cells for 24 hrs, with the addition of the small molecule inhibitors at defined time points. Disruption of the BUNV infection cycle was assessed dually at the 24 hpi time point; firstly, by western blot analysis to quantify rBUNV-WT NP expression and secondly, by quantification of the number of fluorescent rBUNV-eGFP expressing cells compared to solvent controls. The endosomal acidification inhibitor, ammonium chloride (NH_4_Cl), was used as a positive control to delineate the time of virus endosomal escape. Addition of NH_4_Cl to virus infected cells between 0-1 hpi potently inhibited BUNV-specific gene expression (NP and eGFP) compared to addition at the post-entry 10 hpi time point, confirming that endosomal acidification was required for virus internalisation (Fig 3D-E). When NS8593 and clotrimazole were used here, they were most potent when added between 0-2 hpi, causing significant reductions in NP or eGFP expression (Fig 3D, F-G), mirroring the effect of NH_4_Cl and suggesting a role for the corresponding channels during viral entry.

To further corroborate this, we used immunofluorescence microscopy to directly visualise BUNV NP at pre- and post-entry time points (2 and 6 hpi), as determined by the time-of-addition assays (Fig 3H). In the solvent (DMSO)-treated control cells, small distinct puncta of BUNV NP were observed at 2 hpi indicative of incoming virions. By 6 hpi, BUNV NP had flooded the cytoplasm, suggesting that the virus had escaped the endosomal pathway and initiated viral gene expression. Conversely, when endosomal acidification was prevented by the addition of NH_4_Cl, BUNV NP cytoplasmic flooding did not occur, and instead punctate staining was observed at 6 hpi in a similar pattern to that seen at 2 hpi. This indicated no additional BUNV NP expression had occurred, consistent with a block in endosome escape. A similar phenotype was observed for NS8593 and clotrimazole, further supporting a role for *KCNN2* (K_Ca_2.2), *KCNN3* (K_Ca_2.3), *KCNN4* (K_Ca_3.1), *KCNMA1* (BK_Ca_) during the virus entry process.

### Specific *KCNN4* (KCa3.1) inhibitors do not prevent BUNV endosomal escape, despite effect on early infection

As clotrimazole has been shown to inhibit both *KCNN4* (K_Ca_3.1) and *KCNMA1* (BK_Ca_) (Ishii et al., 1997; Rittenhouse et al., 1997), more specific inhibitors of *KCNN4* were sought to delineate the effects of the two channels upon BUNV; namely NS6180 (Strøbæk et al., 2012) and TRAM34 (Wulff et al., 2000) (Fig 4A). At the highest non-toxic concentrations, treatment with NS6180 (30 µM) or TRAM34 (50 µM) in the drug assays described above resulted in a reduction in the number of rBUNV-eGFP infected cells to 40% and 14% of the solvent controls, respectively (Fig 4B-C). Furthermore, in the time-of-addition assays, both drugs were most potent against BUNV when added between 0-2 hpi, significantly reducing both rBUNV-WT NP expression and rBUNV-eGFP expression compared to addition at 10 hpi (Fig 4D-F). These data supported a role for *KCNN4* (K_Ca_3.1) during early infection.

**Figure 4.**
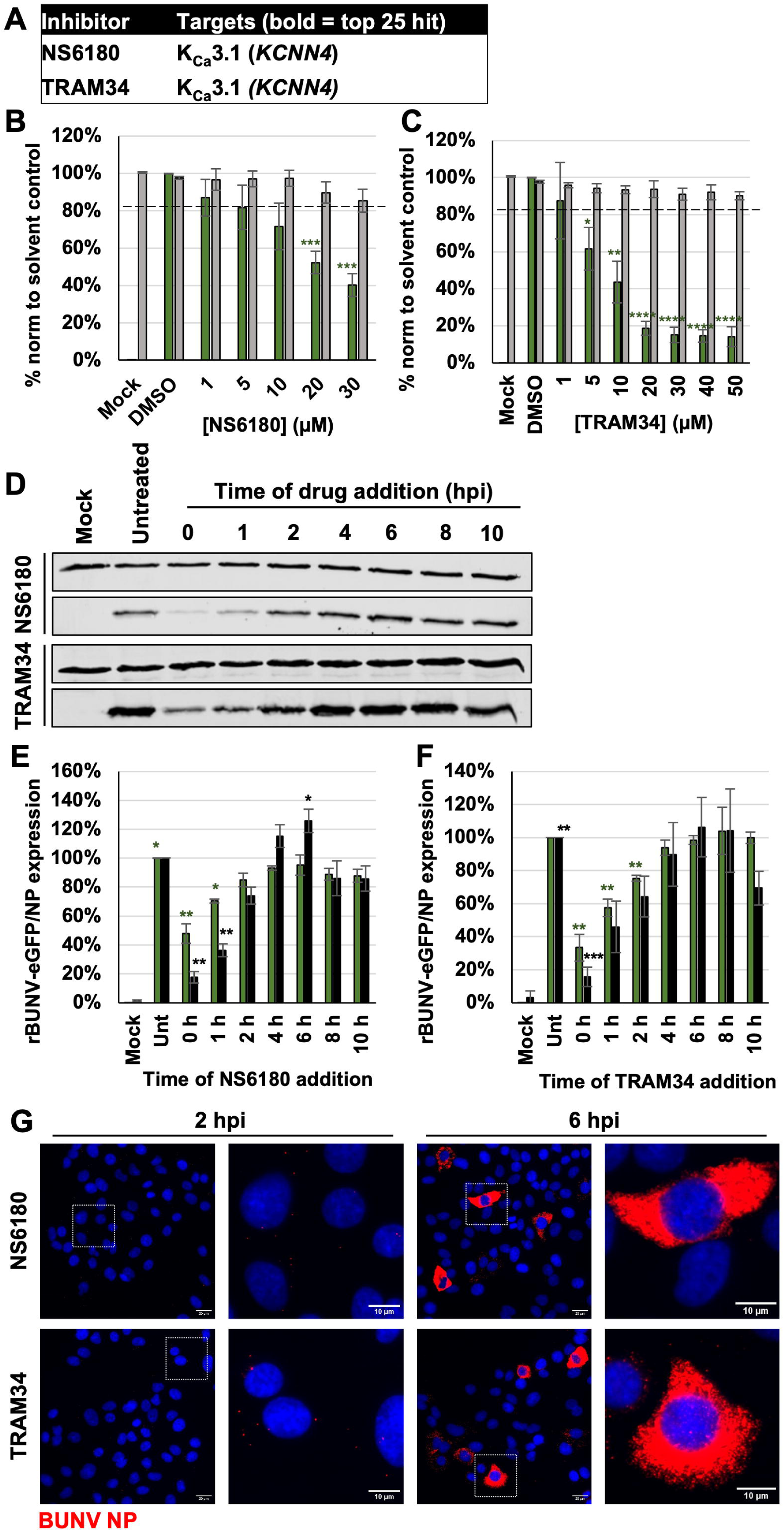
*KCNN4* (K_Ca_3.1) plays a role in early infection but not in endosomal escape. A. Table to show targets of NS6180 and TRAM34. B-C. A549 cells were pre-treated with small molecule inhibitors in A. After 45 mins of treatment, cells were infected with rBUNV-eGFP at MOI 0.1 for 24 hours. The number of rBUNV-eGFP expressing cells was quantified using IncuCyte S3 live imaging and normalised to the solvent control (green bars). Averages of three biological repeats are shown, each performed in duplicate. Drug cytotoxicity was measured immediately afterward via MTS assay and normalised to solvent controls (grey bars), The dashed line represents the toxicity threshold. Three biological repeats were performed, each in triplicate. D-F. A549 cells were infected with either rBUNV-WT or rBUNV-eGFP at MOI 0.5 for 24 hours. At the indicated time points post-infection, infected cells were treated with the NS6180 or TRAM34. Cells in the 0 hpi condition were pre-treated for 45 mins prior to infection. D. Representative western blot analysis of BUNV NP and GAPDH expression. E-F. Densitometry analysis of rBUNV-WT NP expression (black bars) alongside number of rBUNV-eGFP expressing cells quantified by IncuCyte S3 (green bars), normalised to untreated (solvent) controls. Data is averaged from three biological repeats. One way ANOVA tests were performed to assess statistical significance. P< 0.05*, P<0.01**, P<0.001***, P<0.0001****. G. A549 cells were pre-treated with NS6180 or TRAM34 for 45 mins. Cells were incubated on ice and rBUNV-WT was added at MOI 5 for 1 hour to promote virus binding but not internalisation. Non-internalised virus was removed, and infection was allowed to progress in the presence of the inhibitor at 37°C. Cells were fixed at 2 or 6 hpi and immuno-stained for BUNV NP (red) and DAPI (blue). Images were taken on Olympus IX83 widefield deconvolution microscope at 60x magnification. White boxes indicate zoomed field of view. Scale bars are included (20 µM wide, 10 µM zoom).

However, in BUNV-infected cells pre-treated with NS6180 or TRAM34, NP flooded the cytoplasm at 6 hpi (Fig 4G), displaying the same phenotype as the solvent control (Fig 3H). As viral gene expression was occurring in these treated cells, this indicated that inhibition of *KCNN4* (K_Ca_3.1) alone was not sufficient to block viral endosomal escape and may play a role in another, as yet unknown, early-stage event.

### Inhibition of select K_IR_ and K_V_ channels also blocks BUNV endosomal escape

While investigating the role of the other K^+^ channel families which appeared on the list of the 25 highest ranked K^+^ channels, we were limited by the availability of specific small molecule inhibitors. Several inwardly rectifying K^+^ channel hits were targeted by our panel of inhibitors (Fig 5A) with VU590 blocking *KCNJ1* (ROMK/K_IR_1.1) and *KCNJ13* (K_IR_7.1) (Lewis et al., 2009), and ML133 acting upon *KCNJ2* (IRK1/K_IR_2.1) (H. R. Wang et al., 2011). Treatment using these compounds resulted in a decrease in the number of rBUNV-eGFP infected cells to 51% and 24% of the solvent controls, respectively (Figs 5B-C). Lastly, voltage-gated channel *KCNQ2* (K_V_7.2/K_V_LQT2) was blocked by the action of ML252 (40 µM) (Cheung et al., 2012), resulting in a reduction of rBUNV-eGFP infection to 36% of the solvent control (Fig 5D). Taken together, these findings corroborate the outcome of the siRNA screen and confirm the importance of multiple K^+^ channels during BUNV infection.

**Figure 5.**
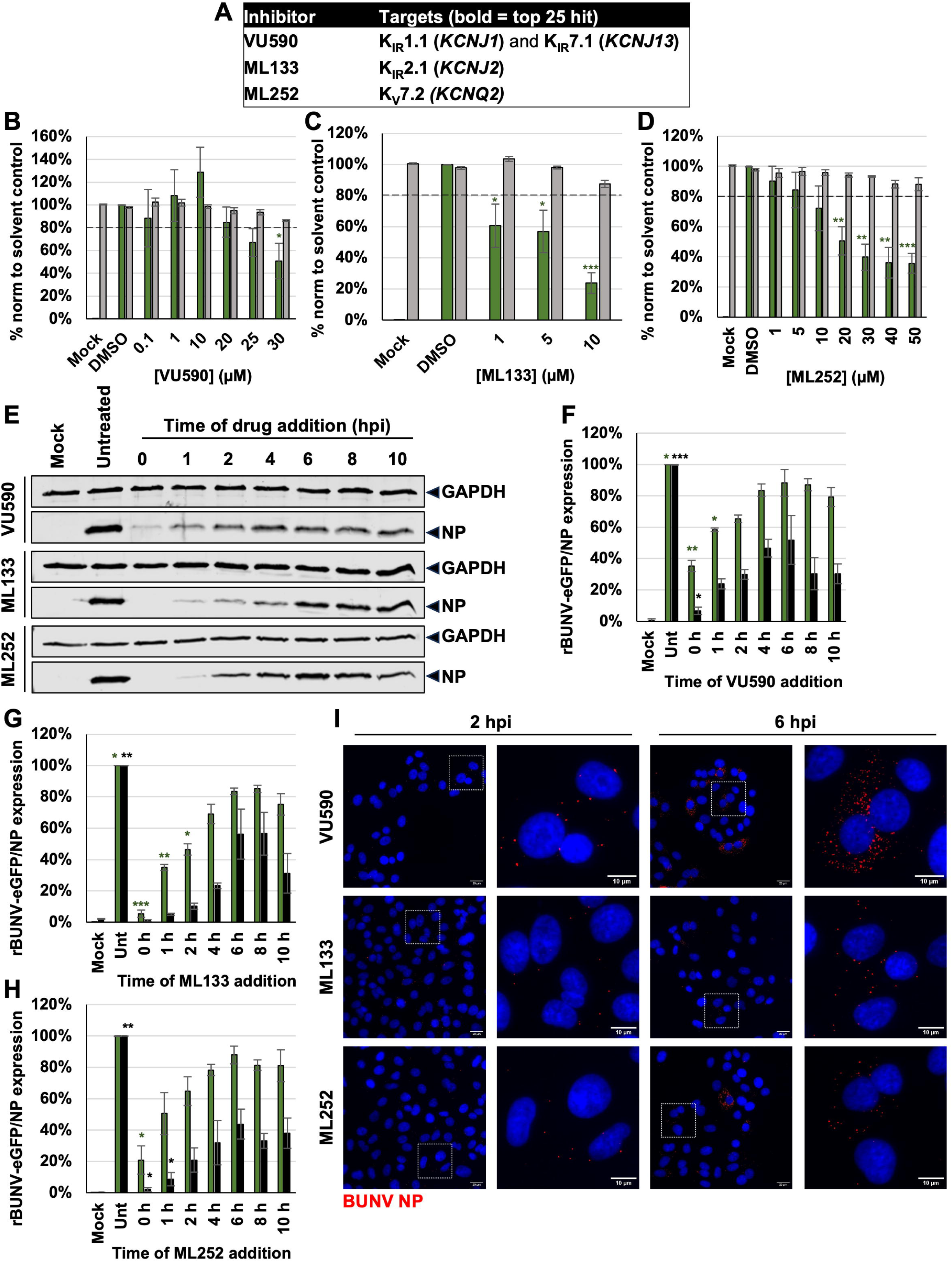
Inhibition of select K_IR_ and K_V_ channels blocks BUNV endosomal escape. A. Table to show targets of small molecular inhibitors targeting K^+^ channels. Targets in bold were hits in the siRNA screen. Channel protein names are shown, with gene names in brackets. B-D. A549 cells were pre-treated with small molecule inhibitors in A. After 45 mins of treatment, cells were infected with rBUNV-eGFP at MOI 0.1 for 24 hours. The number of rBUNV-eGFP expressing cells was quantified using IncuCyte S3 live imaging and normalised to the solvent control (green bars). Averages of three biological repeats are shown, each performed in duplicate. Drug cytotoxicity was measured immediately afterward via MTS assay and normalised to solvent controls (grey bars), The dashed line represents the toxicity threshold. Three biological repeats were performed, each in triplicate. E-H. A549 cells were infected with either rBUNV-WT or rBUNV-eGFP at MOI 0.5 for 24 hours. At the indicated time points post-infection, infected cells were treated with inhibitors in A. Cells in the 0 hpi condition were pre-treated for 45 mins prior to infection. Representative western blot analysis of BUNV NP and GAPDH expression is shown in E. Graphs F-H show the densitometry analysis of rBUNV-WT NP expression (black bars) alongside number of rBUNV-eGFP expressing cells quantified by IncuCyte S3 (green bars), normalised to untreated (solvent) controls. Data is averaged from three biological repeats. One way ANOVA tests were performed to assess statistical significance. P< 0.05*, P<0.01**, P<0.001***, P<0.0001****. I. A549 cells were pre-treated with the inhibitors in A for 45 mins. Cells were incubated on ice and rBUNV-WT was added at MOI 5 for 1 hour to promote virus binding but not internalisation. Non-internalised virus was removed, and infection was allowed to progress in the presence of the inhibitor at 37°C. Cells were fixed at 2 or 6 hpi and immuno-stained for BUNV NP (red) and DAPI (blue). Images were taken on Olympus IX83 widefield deconvolution microscope at 60x magnification. White boxes indicate zoomed field of view. Scale bars are included (20 µM wide, 10 µM zoom).

Time-of-addition assays were employed to determine the stage of infection at which these channels were required, and it was revealed that VU590 and ML252 were most potent against rBUNV-eGFP when added between 0-1 hpi (Fig 5E, F, H), and ML133 significantly inhibited infection up to 2 hpi (Fig 5G), similar to NH_4_Cl and K_Ca_ inhibitors described above. Furthermore, studies into BUNV NP localisation upon treatment with these inhibitors also showed the same phenotype as NH_4_Cl and the K_Ca_ inhibitors, with distinct NP puncta observed at 6 hpi (Fig 5I). Taken together, blockade of *KCNJ1* (K_IR_1.1), *KCNJ13* (K_IR_7.1), *KCNJ2* (K_IR_2.1) and *KCNQ2* (K_V_7.2) prevented BUNV endosomal escape during early infection.

Of note, when using ammonium chloride, NS8593, clotrimazole, VU590, ML133 and ML252 at later time points, a full recovery of NP expression to DMSO-only levels was not observed, likely due to their inhibitory effects on virus entry precluding a subsequent round of BUNV multiplication within the 24 hr analysis window. In contrast, late time point treatment with compounds NS6180 and TRAM34 (*KCNN4* inhibitors) resulted in near recovery of BUNV NP levels, likely due to these compounds having a lesser effect on virus entry and therefore allowing a second round of virus entry/infection during the 24 hr window, and thus increased BUNV NP accumulation. This phenomenon was not observed with rBUNV-eGFP, likely due to the stability of eGFP expression within infected cells.

### BUNV requires K^+^ channels which localise to late endosomes

Since BUNV enters cells by endocytosis and our data suggested that K_Ca_s (*KCNN2, KCNN3, KCNMA1*), K_IR_s (*KCNJ1, KCNJ2, KCNJ13*) and K_V_s (*KCNQ2*) played a role in virus entry, we hypothesized the channel proteins would localise to either early endosome (EE) or LE compartments. To test this, the cellular localisation of these channels was next investigated using confocal microscopy. To achieve this, we generated cDNAs to express the 7 selected channels as wildtype mCherry fusion proteins in A549 cells, allowing visualisation of channel location in relation to a panel of endosomal markers, namely EEA1 (EE) and CD63 or Rab7 (LE).

Interestingly, no co-localisation was observed between any of the K^+^ channels and EEA1, indicating an absence of expression of these channels in EEs (Supp Fig 3). However, co-localisation was observed between CD63 and K_IR_1.1 (*KCNJ1*), K_IR_2.1 (*KCNJ2*), K_IR_7.1 (*KCNJ13*) and BK_Ca_ (*KCNMA1*) (Fig 6), and these results were confirmed by co-localisation with LE marker Rab7 (Supp Fig 4). Taken together, these data confirm the presence of three K_IR_s and one K_Ca_ channels at LEs and support that the notion that these channels contribute to the high [K^+^] required for BUNV fusion with endosomes and the subsequent release of its genomic segments into the cytoplasm.

**Figure 6.**
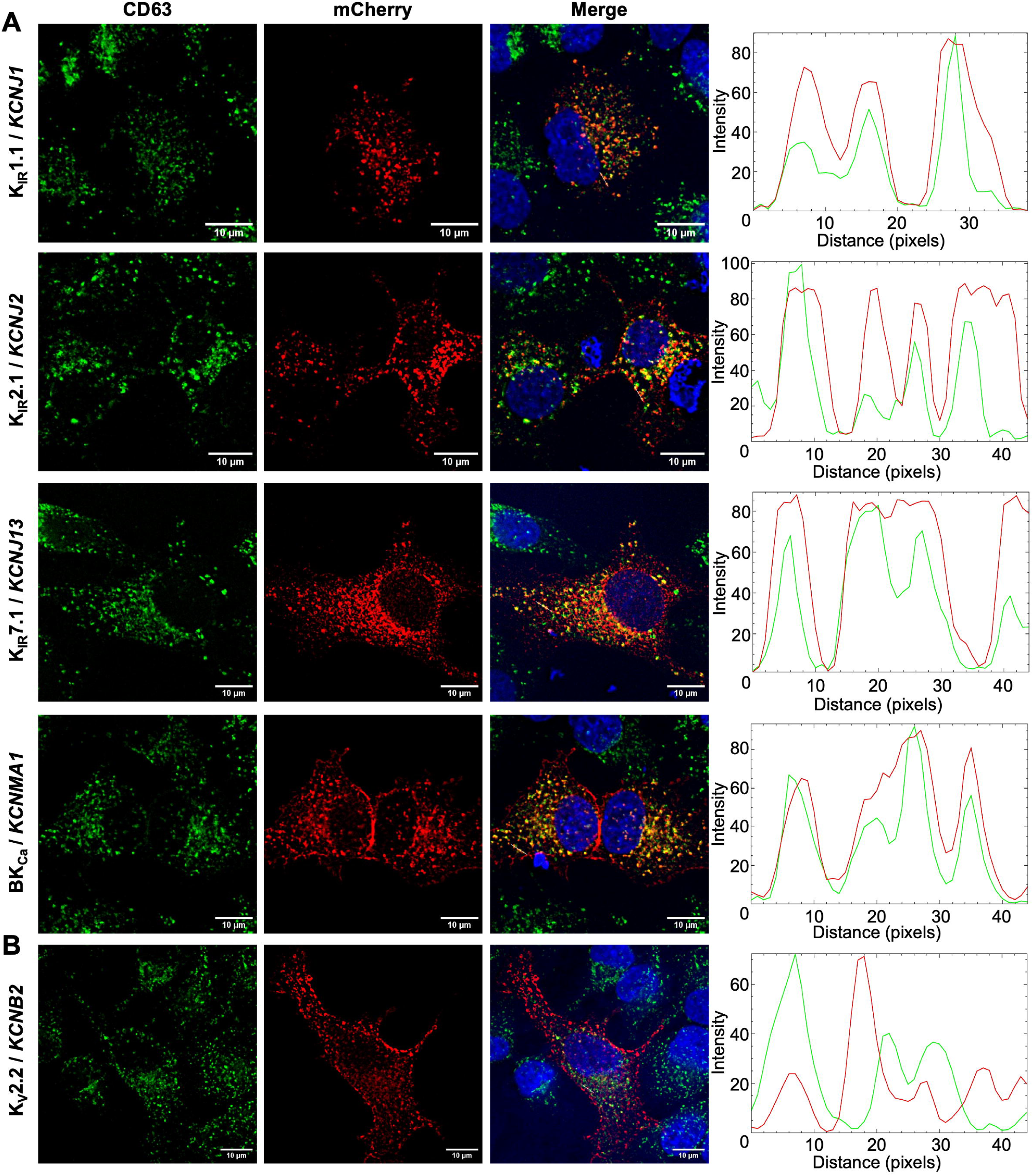
K^+^ channels co-localise with late endosomal marker CD63. Plasmids encoding WT K^+^ channels tagged with mCherry (red) were transfected into A549 cells for 48 hours. Cells were then fixed and immuno-stained for late endosomal marker CD63 (green). DAPI is also shown (blue). Cells were imaged on Olympus IX83 widefield deconvolution microscope at 60x magnification. A. Channels which co-localise to LE. B. Negative control. Scale bars (10 µM) and line scan analysis performed on FIJI software.

As a negative control, we selected *KCNB2* based on a lack of impact on viral infection during the siRNA screen (Supp Fig 1). As expected, K_V_2.2 (*KCNB2*) did not show any co-localisation with any endosomal markers and showed cytoplasmic and cell membrane distribution, further confirming the specificity of our channels of interest. Intriguingly, the remaining three channels of interest, namely K_Ca_2.2 (*KCNN2*), K_Ca_2.3 (*KCNN3*) and K_V_7.2 (*KCNQ2*), shown above to be involved in BUNV entry, failed to co-localise with CD63 or Rab7 (Fig 7, Supp Fig 4), and instead displayed a punctate cytoplasmic localisation. This finding suggests that while K_Ca_2.2 (*KCNN2*), K_Ca_2.3 (*KCNN3*) and K_V_7.2 (*KCNQ2*) play important roles in the BUNV entry process, they likely act at a stage distinct from endosomal escape.

**Figure 7.**
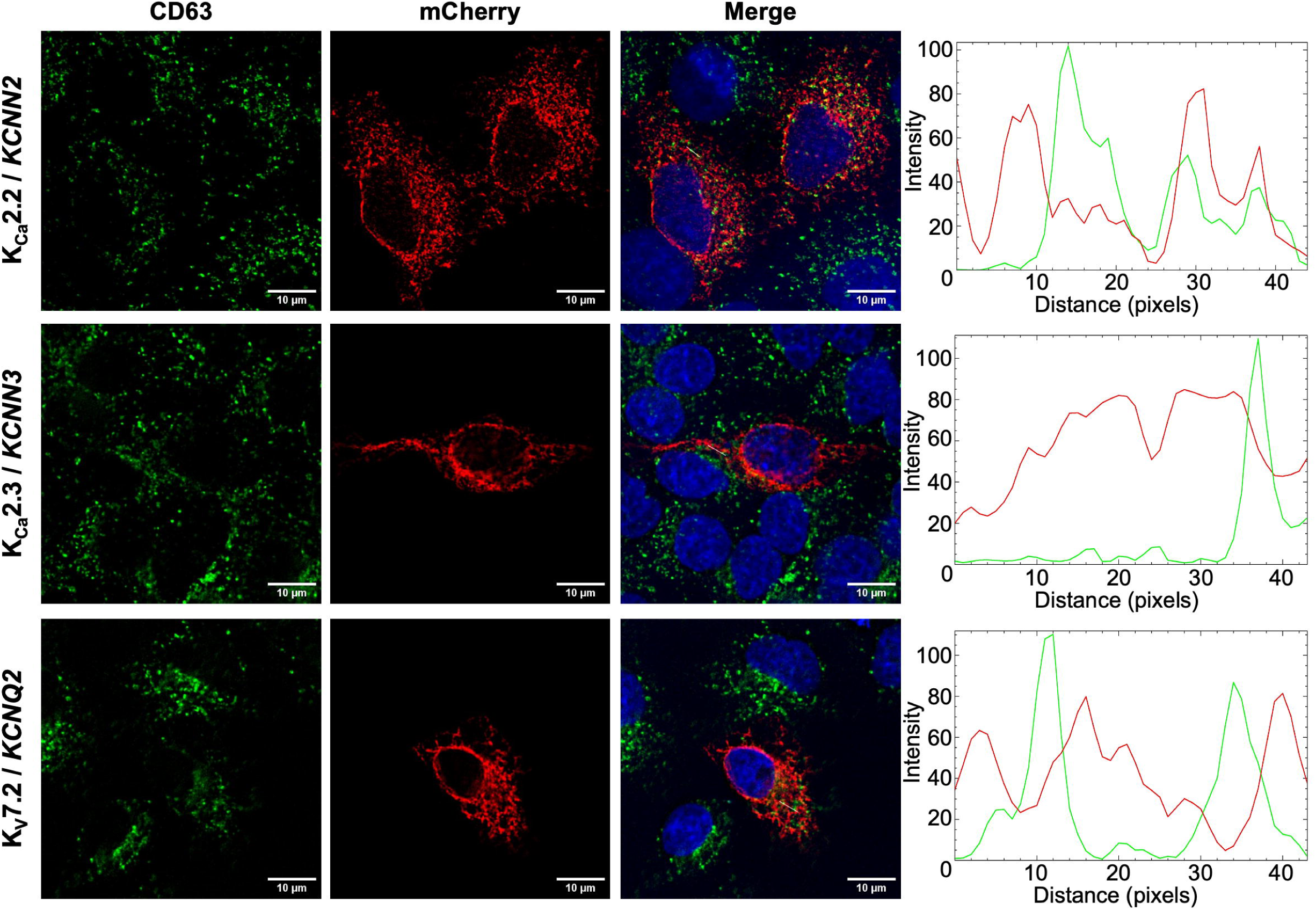
KCNN2, KCNN3 and KCNQ2 K^+^ channels do not co-localise with late endosomal marker CD63. Plasmids encoding WT K^+^ channels tagged with mCherry (red) were transfected into A549 cells for 48 hours. Cells were then fixed and immuno-stained for late endosomal marker CD63 (green). DAPI is also shown (blue). Cells were imaged on Olympus IX83 widefield deconvolution microscope at 60x magnification. Scale bars (10 µM) and line scan analysis performed on FIJI software.

### Generation of non-functional K^+^ channel pore mutants decreases BUNV infection

To further investigate the importance of the 7 K^+^ channels of interest during BUNV entry, and to avoid any off-target effects of the small molecule inhibitors, we used an alternative approach to block K^+^ channel activity. K^+^ channels are highly selective, with ion selectivity being determined by a TxGYG motif that lies within the channel pore (Doyle et al., 1998; Zhou et al., 2001) and mutation of this motif renders the channel unable to transport K^+^ ions (Dahal et al., 2012; Preisig-Müller et al., 2002). This creates a dominant-negative effect wherein co-assembly of wildtype and mutant K^+^ channel subunits inhibits the pore function (Lafrenière et al., 2010; Reed et al., 2016; Zerr et al., 1998). Thus, we created dominant-negative, non-conducting, mCherry-fused channel subunits by mutation of the GYG motif to AAA in each subunit, followed by assessment of how expression of each channel mutant affected rBUNV-eGFP multiplication.

Interestingly, transfection of A549 cells with the channel subunit-expressing cDNAs rendered these cells poorly infectable with rBUNV-eGFP, likely due to detection of the transfected plasmids by the cGAS-STING innate immune pathway (Langereis et al., 2015). Therefore, we instead used HEK-293T cells for this experiment, which lack an interferon response. To validate the use of this alternative cell line, the expression of our 7 K^+^ channels of interest was verified by RT-qPCR (Supp Fig 2B).

Following transfection of wildtype (WT) or pore mutant (AAA) K^+^ channel cDNAs, cells were infected with rBUNV-eGFP, and at 24 hpi, cells were harvested and analysed by flow cytometry. Cells positive for mCherry fluorescence, indicating expression of exogenous channels, were selected and the intensity of green fluorescence was measured as an indication of rBUNV-eGFP infection levels. For the non-endosomal channel, *KCNB2* (K_V_2.2), rBUNV-eGFP expression was not significantly altered when either exogenous *KCNB2*-WT or non-functional *KCNB2*-AAA was expressed in cells (Fig 8), confirming that this channel was not involved in virus infection.

**Figure 8.**
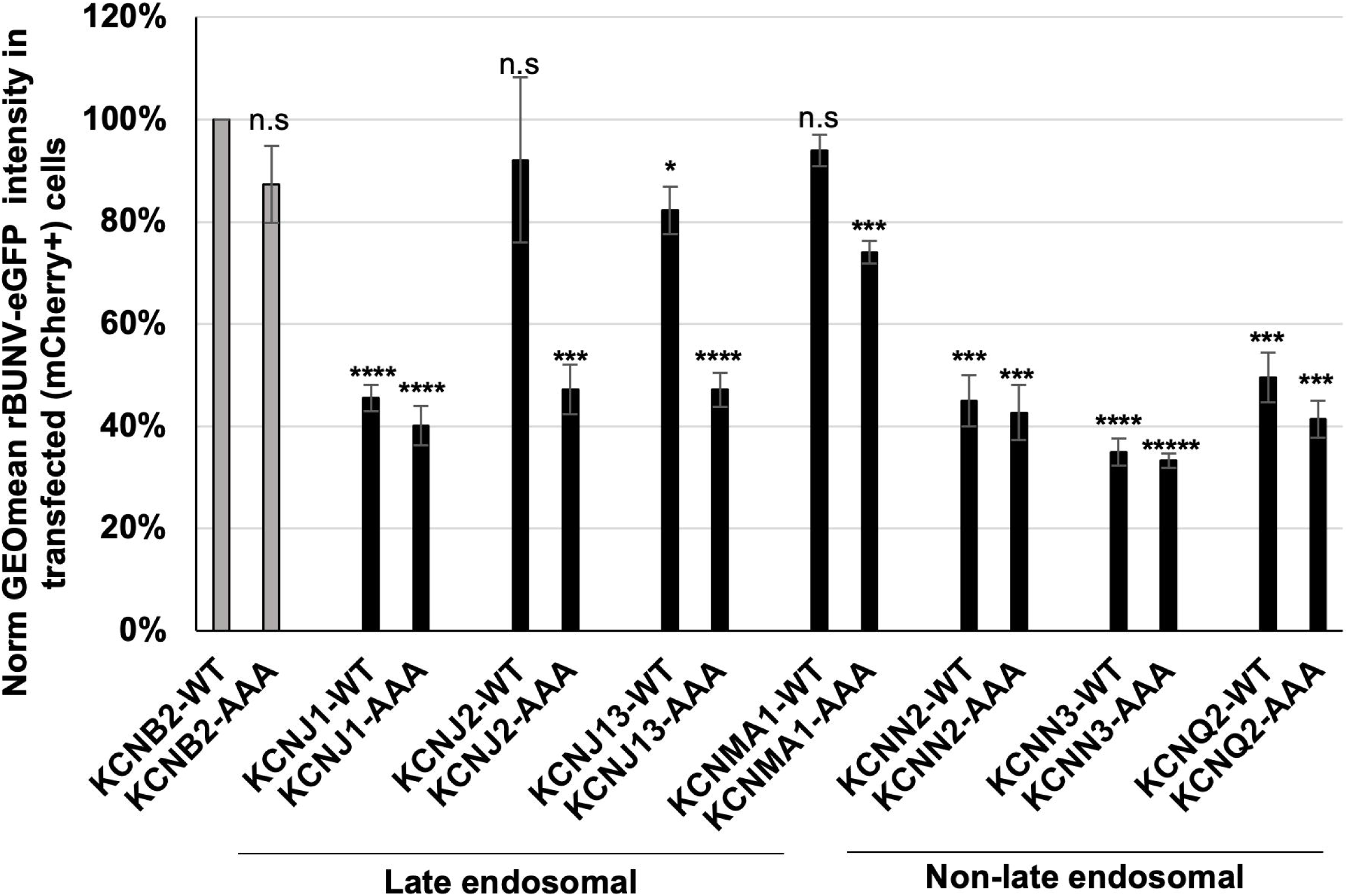
Generation of non-functional K^+^ channel pore mutants decreases BUNV infection. Plasmids encoding WT or dominant-negative, non-functional pore mutant (AAA) K^+^ channels tagged with mCherry were transfected into HEK-293T cells. After 24 hours, cells were infected with rBUNV-eGFP at MOI 2 for a further 24 hours. Cells were harvested for FACS analysis. For a population of >10,000 mCherry-positive cells the GEOmean of eGFP signal was measured. Values were normalised to cells transfected with KCNB2-WT as the negative control (grey), and the average of three biological repeats is shown. Error bars represent standard error. P< 0.05*, P<0.01**, P<0.001***, P<0.0001****.

The non-functional pore-mutants showed a statistically significant reduction rBUNV-eGFP expression for three of the four channels found to co-localise at LE; *KCNJ2*-AAA (47%), *KCNJ13*-AAA (47%) and *KCNMA1*-AAA (74%), when compared to *KCNB2*-WT. Furthermore, the expression of *KCNJ13*-WT also had a statistically significant impact on infection, although this was to a lesser extent than the mutant (82%). However, despite also being found in LE and having a role during virus entry, expression of *KCNJ1*-AAA did not affect rBUNV-eGFP expression any more significantly than that of overexpressing *KCNJ1*-WT (40% vs 46%, respectively).

We also studied the three K^+^ channels that did not co-localise to any endosomal compartments but did show an effect on virus entry: *KCNN2* (K_Ca_2.2), *KCNN3* (K_Ca_2.3), and *KCNQ2* (K_V_7.2). For these channels, expressing either the WT or pore-mutant impacted rBUNV-eGFP expression similarly, and all were <50% of the negative control *KCNB2* (K_V_2.2).

Despite this unexpected consequence of WT channel expression, the expression of AAA pore mutants of *KCNJ2* (K_IR_2.1), *KCNJ13* (K_IR_7.1) and *KCNMA1* (BK_Ca_) resulted in a decrease in rBUNV-eGFP expression which was more significant than the corresponding WT when compared to the negative control. Taking these results together with the endosomal location of these three channels, and the arrest of virion escape following channel inhibition, our results support a critical role for K_IR_2.1 (*KCNJ2*), K_IR_7.1 (*KCNJ13*) and BK_Ca_ (*KCNMA1*) in the endosomal stages of BUNV entry.

## Discussion

Beyond the well-established pH gradient, the ionic milieu of the endocytic network is complex and controlled by multiple ion channels, transporters and exchangers. For K^+^, the concentration rises rapidly from extracellular levels in EE after endocytosis and continues to rise through LE and lysosomes (Scott & Gruenberg, 2011), however the functional roles of this high endosomal [K^+^] are not well understood. Following on from our previous work which highlighted the importance of high endosomal [K^+^] during bunyavirus entry (Hover et al., 2016, 2018, 2023), here we aimed to identify the ion channel(s) responsible for this accumulation of ions.

We have demonstrated that a wide range of K^+^ channels from different K^+^ channel families play a role in the entry of BUNV into cells. Through a comprehensive siRNA library targeting human ion channels, we found that the knockdown of 19 individual K^+^ channels inhibited BUNV infection by >50%, potentially indicating multiple channels exert a combinatorial effect over K^+^ flux in relation to BUNV infection.

Focussing on channels for which small molecule inhibitors were available, we performed time-of-addition assays to identify K^+^ channels which played a role during the entry stage of infection. Of note, when using ammonium chloride, NS8593, clotrimazole, VU590, ML133 and ML252 at later time points, a full recovery of NP expression to DMSO-only levels was not observed, likely due to their inhibitory effects on virus entry precluding a subsequent round of BUNV multiplication within the 24 hr analysis window. In contrast, late time point treatment with compounds NS6180 and TRAM34 (*KCNN4* inhibitors) resulted in near recovery of BUNV NP levels, likely due to these compounds having a lesser effect on virus entry and therefore allowing a second round of virus entry/infection during the 24 hr window and thus increased BUNV NP accumulation. This phenomenon was not observed with rBUNV-eGFP, likely due to the stability of eGFP expression within infected cells.

Using confocal microscopy, we were able to show a blockade of BUNV endosomal escape in the presence of K^+^ channels inhibitors NS8593, clotrimazole, VU590, ML133 and ML252, strongly suggesting the channels that are susceptible to these drugs reside within endosomal compartments. To confirm this, we showed that four of those drug-susceptible channels were localised to LE; namely K_IR_1.1 (*KCNJ1*), K_IR_2.1 (*KCNJ2*), K_IR_7.1 (*KCNJ13*) and BK_Ca_ (*KCNMA1)*. Expressing non-functional K_IR_2.1 (*KCNJ2*), K_IR_7.1 (*KCNJ13*) and BK_Ca_ (*KCNMA1)* pore mutant channels significantly decreased BUNV infection, and these observations together suggest a functional role for these channels in maintaining the ionic balance of the LE which is the site of BUNV fusion and escape into the cytoplasm.

The BK_Ca_ (*KCNMA1*) channel allows the passive flow of K^+^ ions along the electrochemical gradient into endosomes. It is activated by both Ca^2+^- and voltage and is characterised by a large K^+^ conductance (Kendall & Holian, 2024). Co-localisation between BK_Ca_ and lysosomal protein LAMP-1 was observed previously (Cao et al., 2015; W. Wang et al., 2017), while no-colocalisation was observed with EEA1 (W. Wang et al., 2017), similar to our results. Within lysosomes, high [Ca^2+^] was shown to activate BK_Ca_ resulting in an influx of K^+^ as a counterion for the release of Ca^2+^ into the cytoplasm via TRPML1 (Cao et al., 2015). Therefore, as BK_Ca_ and TRPML1 both co-localise with LAMP-1 which is also found in LE compartments (Dong et al., 2010) containing a high [Ca^2+^], we hypothesize that the activation of BK_Ca_ channels could also occur within LEs, providing the high [K^+^] that is required for bunyaviral fusion and endosomal escape.

The protein products of *KCNJ1*, *KCNJ2* and *KCNJ13* form tetrameric K_IR_1.1, 2.1 and 7.1 channels, respectively (Lopatin et al., 1994; Matsuda et al., 1987). Although they are known to be internalised by clathrin-dependent endocytosis (Hibino et al., 2010), we are the first to propose a functional role for these channels within LE. *KCNJ1* (K_IR_1.1) is highly sensitive to cytoplasmic pH, being inhibited below pH 6.8 (Choe et al., 1997), therefore it is likely that the *KCNJ1* (K_IR_1.1) channels we observed at LE (pH 5.5) would be actively moving K^+^ ions and contributing to the accumulation of K^+^ within LE. Whilst part of the same family, *KCNJ2* (K_IR_2.1) and *KCNJ13* (K_IR_7.1) are only weakly-sensitive to pH (Hibino et al., 2010) and the single channel conductance of *KCNJ13* (K_IR_7.1) is very small compared to the other K_IR_ channels (Krapivinsky et al., 1998). These differences perhaps allow for further fine tuning of ionic balances within LE.

Although *KCNN2*, *KCNN3* and *KCNQ2* were also susceptible to the compounds which inhibited BUNV entry in time-of-addition assays, we did not detect their expression in EE or LE. Overexpression of these WT channels also significantly decreased BUNV infection, however there was no significant difference in infection between overexpression of WT channels or expression of dominant negative pore mutants for these three channels. *KCNN2* and *KCNN3* encode subunits of the tetrameric SK2/K_Ca_2.2 and SK3/K_Ca_2.3 channels respectively. *KCNN2* (K_Ca_2.2) and *KCNN3* (K_Ca_2.3) are voltage-independent, Ca^2+^-activated channels. Meanwhile *KCNQ2* encodes a voltage- and lipid-gated channels protein known as K_v_7.2/K_v_LQT2. It is unclear how blockade of the non-endosomal channels was able to affect virus entry, perhaps they contribute to the ionic balance of LE compartments indirectly, or perhaps they are trafficked/recycled/internalised through endosomal compartments wherein their activity contributes to the endosomal [K^+^], but their expression level is too low to be visualised by these methods. There is still a lot more to be learned about the interplay between these channels and how K^+^ homeostasis is maintained within cells, and endosomes in particular.

We suggest that a broad range of K^+^ channels co-ordinate together to regulate K^+^ accumulation within LE, and that this is disrupted when one or more of these channels are inhibited. The consequences of this disruption on cellular proteins currently remains unknown, and further studies into this may elucidate the native purpose of the K^+^ gradient within the endocytic network. As we were only able to target a subset of the K^+^ channel hits from the siRNA for further investigations, it is highly likely that more of the top 25 K^+^ channel hits from the siRNA screen also contribute to the ionic balance within the endocytic network. The three K^+^ channels we have highlighted here (*KCNJ2*/K_IR_2.1, *KCNJ13*/K_IR_7.1 and *KCNMA1*/BK_Ca_) contribute to endosomal K^+^ accumulation and therefore represent novel host factors for anti-bunyaviral therapeutics.

## Materials and Methods

### Cell lines

A549, BHK-21, BSR-T7 and HEK293T cells were acquired from American Type Culture Collection and maintained in high-glucose Dulbecco’s modified Eagle medium (DMEM, Sigma-Aldrich), supplemented with 10% heat-inactivated fetal bovine serum (FBS), 100 µg of streptomycin/mL, and 100 U of penicillin/mL (P/S), and incubated in a humidified incubator at 37°C with 5% CO_2._ Additionally, BSR-T7 cells constitutively expressing T7 RNA polymerase (T7P) were supplemented with G418 (500 µg/mL) every other passage.

### Plasmid design

Rescue plasmids encoding full-length BUNV antigenome-sense BUNV RNA transcripts: pT7riboBUNS(+), pT7riboBUNM(+), and pT7riboBUNL(+) have been previously described (Bridgen & Elliott, 1996). Plasmid pT7riboBUNS-eGFP(+) was generated as recently described for rLCMV-eGFP (Shaw et al., 2024). Briefly, the eGFP-porcine teschovirus 1 2A (P2A) open reading frame (ORF) was subcloned into pT7riboBUNS(+) using complementary flanking restriction sites. The eGFP-P2A ORF was inserted between the 5’ untranslated region (UTR) and the NP ORF start codon and the self-cleaving P2A peptide sequence (ATNFSLLKQAGDVEENPGP) separated eGFP and NP ORFs, allowing independent expression of eGFP and NP upon infection.

### Recovery of recombinant BUNV-eGFP

BSR-T7 cells were seeded at 2×10^5^ cells per well in a 6-well plate. Cells were transfected with 1 µg pT7riboBUNS-eGFP(+), 1 µg pT7riboBUNM(+), and 1 µg pT7riboBUNL(+), alongside 0.3 µg of a T7P support plasmid in OptiMEM (Gibco) uwing 2.5 µL/µg TransIT-LT1 transfection reagent (Mirus). After 4 hrs incubation at 33°C, the transfection mix was removed and replaced with DMEM containing 2.5% FBS and 1% P/S. At 5 days post-transfection (dpt), the supernatant was clarified by centrifugation and transferred to fresh BHK-21 cells in a T25 flask. The BSR-T7 cells were also lysed to check for viral protein production by western blotting. The BHK-21 cells were incubated at 37°C for 2-3 days whilst being monitored for green fluorescence and cytopathic effect (CPE). Viral supernatant was collected, clarified and filtered before being aliquoted and snap-frozen for storage at −80°C. BHK-21 cell lysates were also collected for western blotting analysis.

### Virus titration

Virus was titrated by plaque assay, wherein a confluent monolayer of BHK-21 cells in a 12-well plate were infected with serial dilutions of viral supernatant in serum-free DMEM for 1 hr at 37°C. Inoculum was removed and replaced with a 1.6% methylcellulose overlay diluted 1:1 in DMEM containing 2.5% FBS and 1% P/S. Plaque assays were incubated at 37°C for 3 days without moving, after which the overlay was removed, the cells were washed in 1x phosphate buffered saline (PBS) and fixed by incubation with 4% paraformaldehyde (PFA) on ice for 10 mins. For BUNV-WT, plaques were visualised by staining with 2% crystal violet in 20% ethanol, and for rBUNV-eGFP, fluorescent plaques were visualised by whole-well scanning on the IncuCyte S3 live imaging platform (Sartorius).

### Reverse transfection of siRNA library

The Silencer™ Human Ion Channel siRNA Library (Ambion) was used as described previously (Shaw et al., 2024). In each well of a 96-well plate 0.3 µL of Lipofectamine RNAiMAX transfection reagent (Thermo) was added to 16.7 uL Opti-MEM. A volume of 3 uL siRNA was then pipetted into this to a concentration of 3 pmol and incubated at room temperature for 20 mins. Scrambled siRNA controls were utilised, alongside controls containing only the transfection reagent. A549 cells were trypsinised and resuspended in Opti-MEM, and 1.25×10^4^ cells were added per well and incubated at 37°C for 24 hrs. The transfection media was gently removed and replaced with DMEM containing 10% FBS and 1% P/S to allow the cells to recover. At 48 hours post-transfection (hpt), cells were infected with rBUNV-eGFP at MOI 0.5 for a further 24 hrs. The infection was assessed at 72 hpt via IncuCyte S3 quantification of the total integrated intensity of the green (eGFP) fluorescent signal.

### Two-step RT-qPCR

Untreated A549 cells were harvested from a confluent 6 well-plate by trypsinisation and pelleted by centrifugation at 1000 rpm. Cells were resuspended in 1x PBS to wash them before being pelleted again. Pellets were either stored at −80°C or used immediately. The Monarch Total RNA Miniprep Kit (New England Biolabs, NEB) was used to extract the total RNA from A549 cells following the manufacturers protocol. First stand cDNA synthesis of total RNA was performed using LunaScript RT Supermix Kit (NEB) according to the protocol.

Amplification of gene specific DNA was performed using Luna Universal qPCR Master Mix (NEB) with the primers outlined in Table 1. A volume of 4 µL cDNA product was used in the qPCR reaction, and samples were normalised to GAPDH expression. PCR products were run on a 2% agarose gels to confirm correct amplicon size.

**Table 1.**
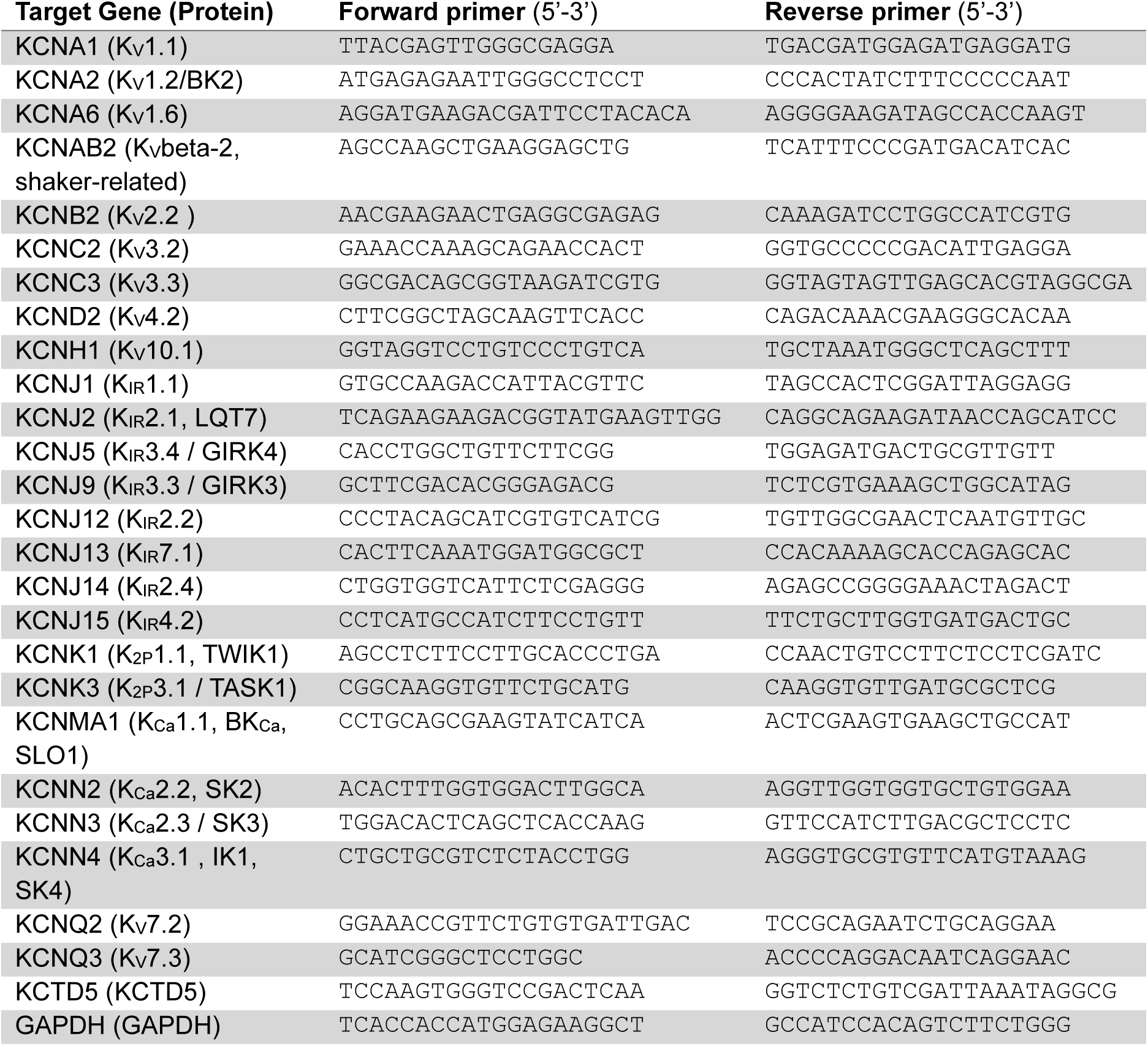
Details of the primers used in qPCR experiments to measure comparative mRNA expression levels of K^+^ channels and GAPDH in A549 cells.

### K^+^ channel inhibitor treatments

A panel of 7 small molecule inhibitors of K^+^ channels (Table 2) were made up to working concentrations in DMEM containing 2.5% FBS and 1% P/S and used to pre-treat A549 cells for 45 mins prior to virus infection. Treatments were performed in duplicate in a 96-well plate. rBUNV-eGFP was used to infect treated cells at MOI 0.1 in the presence of the inhibitor, or solvent control. At 24 hpi, eGFP fluorescence was measured via IncuCyte S3. Immediately following this, infection media was replaced with fresh complete media supplemented with CellTiter 96 Aqueous One Solution Reagent (MTS, Promega) for 1 hr. The absorbance at 490nm was measured as an indication of the toxicity of treated cells compared to untreated controls and a measurement >20% lower than the control was considered cytotoxic.

**Table 2.** Details on the source and known targets of the small molecule inhibitors used against K^+^ channels.

| Inhibitor | Targets | Source |
| --- | --- | --- |
| NS8593 | K<sub>Ca</sub>2.1 (KCNN1), 2.2 (KCNN2), 2.3 (KCNN3) | Sigma, 492031 |
| NS6180 | K<sub>Ca</sub>3.1 (KCNN4) | Stratech, B5740 |
| TRAM34 | K<sub>Ca</sub>3.1 (KCNN4) | Sigma, T6700 |
| Clotrimazole | K<sub>Ca</sub>3.1 (KCNN4), BK<sub>Ca</sub> (KCNMA1), TRPM2 | Sigma, C6019 |
| VU590 dihydrochloride | K<sub>IR</sub>1.1 (KCNJ1) and K<sub>IR</sub>7.1 (KCNJ13) | Alomone labs, V-130 |
| ML133 | K<sub>IR</sub>2.1 (KCNJ2) | Alomone labs, M-275 |
| ML252 | K<sub>V</sub>7.2 (KCNQ2) | Sigma, SML0723 |

### Time-of-addition assays

A549 cells in a 12-well plate were infected with either BUNV-WT or rBUNV-eGFP at MOI 0.5 in infection media for 24 hours. At the indicated time points post-infection (1, 2, 4, 6, 8, 10 hrs), infected cells were treated with small molecule inhibitors of K^+^ channels at 2x working concentration added directly to the infected media so that the cells received a 1x concentration of inhibitor. Cells in the 0 hpi condition were pre-treated for 45 mins prior to infection, and the virus was added directly to the conditioned media. Uninfected (mock) and untreated (solvent only) controls were included. At 24 hpi, the fluorescence of rBUNV-eGFP infected cells were analysed via IncuCyte S3. BUNV-WT infection was analysed by western blotting.

### Western blotting

Cells were lysed in 1x radioimmunoprecipitation assay (RIPA) buffer [50 mM Tris-HCl, pH 7.5, 150 mM NaCl, 1% (vol/vol) NP40 alternative, 0.5% (wt/vol) sodium deoxycholate, and 0.1% (wt/vol) sodium dodecyl sulfate (SDS)], supplemented with 1× cOmplete, Mini, EDTA-free Protease I inhibitor cocktail (Sigma-Aldrich) for 15 min on ice. Sample lysates were collected and resolved on a 12% SDS-PAGE gel before being transferred to a polyvinylidene fluoride membrane (PVDF) membrane using the Trans-Blot turbo (Bio-Rad) system. The membrane was blocked by incubation in Odyssey blocking buffer (Licor) for 1 hr at room temperature. Proteins of interest were probed using primary antibodies made up in 1:4 blocking buffer:PBS and incubated for 1 hr at room temperature or overnight at 4°C. Details of the primary antibodies used are as follows: anti-BUNV-N antisera (in house, 1:5000); anti-GFP [B-2] (Santa Cruz, sc-9996, 1:1000); anti-GAPDH [6C5] (Santa Cruz, sc-32233, 1:7500). Corresponding secondary antibodies were also made up in 1:4 blocking buffer:PBS and incubated for 1 hr at room temperature protected from light. All secondary antibodies were conjugated to an IRDye® 800CW or 680RD (Licor) and used at 1:10,000. Wash steps between each incubation were performed using PBS supplemented with 0.1% Tween-20. The proteins were visualised by their fluorescence using the Licor Odyssey SA imaging system and densitometry was performed using ImageJ.

### Generation of K^+^ channel pore mutants

Plasmids encoding the coding sequences of KCNJ1 (Entrez gene 3758, clone ID 30915211), KCNJ13 (Entrez gene 3769, clone ID 4817527), KCNN2 (Entrez gene 3781, clone ID 40126005), KCNN3 (Entrez gene 3782, clone ID 3619576) and KCNQ2 (Entrez gene 3785, clone ID 3349625) were purchased from Horizon. pCWXPGR-pTF-Kir2.1 encoding KCNJ2 was a gift from Patrick Salmon (Addgene plasmid #114284). The coding sequences of KCNMA1 and KCNB2 were synthesised by Genewiz (Azenta). The vector Zhi3-mCherry was a gift from Sanford Simon (Addgene plasmid #106295).

PCR was used to amplify the desired gene flanked by engineered restriction enzyme sites. Additionally, the stop codons were removed to allow the tagging of the protein of interest with mCherry. Following the restriction digest of the inserts and vector, the two were ligated to produce Zhi3-KCN*XX*-mCherry (wildtype channels). Next, Q5 site-directed mutagenesis (NEB) was used to mutate the GYG motif within each K^+^ channel to AAA. Products were transformed into NEB 5-alpha competent E. coli cells and DNA was isolated using a Maxiprep kit (Qiagen) according to the manufacturers protocol. Whole plasmid sequencing was used to confirm the correct sequence at each step (Plasmid-EZ, Genewiz). Due to a highly repetitive sequence, Zhi3-KCNMA1-AAA-mCherry was synthesised by Genewiz (Azenta).

### Flow cytometry

HEK-293T cells were seeded into 6-wp at a density of 2.5×10^5^ cells/well 18-24 hrs prior to transfection. Zhi3-KCN*XX*-WT/AAA-mCherry plasmid DNA (6 µg) was added to a tube containing 200 µL Opti-MEM, then 12 µL TransIT-LT1 (Mirus) was added and gently mixed. The transfection mix was incubated at room temperature for 30 mins before being added dropwise to the wells containing 1.8 mL Opti-MEM. Cells were incubated with the transfection mix for 4 hrs at 37°C, after which the transfection media was replaced with DMEM containing 10% FBS and 1% P/S. At 24 hpt, transfected cells were infected with rBUNV-eGFP at MOI 2 in DMEM containing 2.5% FBS and 1% P/S for a further 24 hrs. At 48 hpt, red and green fluorescence were confirmed via IncuCyte S3 analysis, and cells were harvested for flow cytometry. To this end, cells were washed with 1x PBS, trypsinised and collected into tubes. The cells were pelleted gently by centrifugation at 2000 rpm for 5 mins. The cell pellet was resuspended in PBS to wash away remaining media and pelleted again. For fixation, cells were next resuspended in cold 4% PFA and incubated on ice for 10 mins. Following a further PBS wash, cells were resuspended in flow cytometry buffer (2mM EDTA, 5% BSA in PBS). The CytoFlex S flow cytometer (Beckman Coulter) was used to analyse the cells. Cells positive for red fluorescence (561_610-20-A) and therefore successfully expressing WT or mutant K^+^ channels were gated upon, and the GEOmean intensity of green signal (488_525-50-A) and the %GFP+ was taken for each sample as a measure of rBUNV-eGFP infection.

### Immunofluorescence and microscopy

A549 cells were transfected with Zhi3-KCN*XX*-mCherry as above. At 48 hpt, cells were fixed by incubation with 4% PFA on ice for 10 mins, followed by permeabilisation in 0.3% triton-X100 for 15 mins at room temperature. Non-specific binding was blocked by incubation with 5% bovine serum albumin (BSA) in 1x PBS at room temperature for 30 mins. Proteins of interest were stained with primary antibodies made up in blocking buffer for 1 hr at room temperature or overnight at 4°C, followed by secondary antibodies in the same manner. Details of these antibodies are as follows: anti-CD63 [MX-49.129.5] (Santa Cruz, sc-5275, 1:100); anti-Rab7 [D95F2] (Cell signalling technology, 9367, 1:100); anti-EEA1 [C45B10] (Cell signalling technology, 3288, 1:100); anti-mCherry [1B3C6] (Proteintech, 68088, 1:500) or anti-mCherry (Proteintech, 26775, 1:500); BUNV-N antisera (in-house, 1:5000); species-specific IgG conjugated to Alexa Fluor®-488, -594 or -647 (Invitrogen, 1:5000). Coverslips were mounted onto slides using ProLong Gold Antifade Mountant with DAPI (Invitrogen) and imaged using Olympus IX83 widefield deconvolution microscope at 60x magnification. ImageJ was used to add scale bars and undertake line scan analysis.

### Statistical analysis

The statistical significance of data was determined by performing one-way ANOVA tests. Significance was deemed when the values were less than or equal to the 0.05 *P* value.

## Funding

This work was supported by BBSRC project grant BB/V007467/1 to HP, JM, JNB, EH and JL and Wellcome trust studentship 102174/B/13/Z to EJAAT.

## Acknowledgements

We acknowledge Wellcome trust equipment grant 221538/Z/20/Z, which supports the use of the IncuCyte live cell imaging platform. The authors thank Dr Ruth Hughes and Dr Sally Boxall of the bioimaging facility, Faculty of Biological Sciences, University of Leeds for their expert assistance and use of the Zeiss LSM880, funded by Wellcome Trust grant WT104918MA.

## Data availability statement

All data generated or analysed during this study are included in this published article (and its Supplementary Information files).

## Author contributions

HMP, JNB, JM, EWH, MS and JDL conceptualized the study, HMP, SH, KP and EJAAT performed the investigation, analysis and data curation. HMP wrote the original draft, JNB, SH, EWH and JDL performed subsequent manuscript reviewing and editing. JNB, JM, EWH and JDL acquired the funding.

## Conflict of interest

The authors declare no conflict of interest.

## Figure legends

**Supplementary Figure 1. Identification of human ion channels required during BUNV infection using siRNA screening.** Table of siRNA ion channel targets which inhibited rBUNV-eGFP expression by over 50% based on eGFP signal across three unique siRNAs. Each siRNA value represents the mean of four experimental repeats, and the median for each target was calculated. Targets are colour-coded by ion selectivity, and the heatmap shows most inhibitory to rBUNV-eGFP expression (red) to least inhibitory (blue). Values for mock cells, BUNV-infected cells treated with the transfection reagent (TFR) only, BUNV-infected cells treated with scrambled siRNA (SCR) are shown as controls, alongside the results for KCNB2 which was chosen as a negative channel control in later experiments due to its lack of impact on rBUNV-eGFP infection.

**Supplementary Figure 2. Expression of K^+^ channels in A549 cells.** The mRNA expression levels of K^+^ channel siRNA targets was measured in A549 cells (A) or HEK293-T cells (B) using RT-qPCR. Total RNA was isolated from cells, and gene specific primers were used to quantify mRNA expression relative to the housekeeping gene, GAPDH. Data is a result of 2 experimental and 2 biological repeats.

**Supplementary Figure 3. K^+^ channels do not co-localise with early endosomal marker EEA1.** Plasmids encoding WT K^+^ channels tagged with mCherry (red) were transfected into A549 cells for 48 hours. Cells were then fixed and immuno-stained for early endosomal marker EEA1 (green). DAPI is also shown (blue). Cells were imaged on Olympus widefield microscope at 60x magnification. Scale bars (10 µM) and line scan analysis performed on FIJI software.

**Supplementary Figure 4. Some K^+^ channels co-localise with late endosomal marker Rab7.** Plasmids encoding WT K^+^ channels tagged with mCherry (red) were transfected into A549 cells for 48 hours. Cells were then fixed and immuno-stained for late endosomal marker Rab7 (green). DAPI is also shown (blue). Cells were imaged on Olympus IX83 widefield deconvolution microscope at 60x magnification. Scale bars (10 µM) and line scan analysis performed on FIJI software.

## Notes

### Competing Interest Statement

The authors have declared no competing interest.

## References

Bridgen, A., & Elliott, R. M. (1996). Rescue of a segmented negative-strand RNA virus entirely from cloned complementary DNAs. Proceedings of the National Academy of Sciences of the United States of America, 93(26), 15400. 10.1073/PNAS.93.26.15400

Cao, Q., Zhong, X. Z., Zou, Y., Zhang, Z., Toro, L., & Dong, X. P. (2015). BK Channels Alleviate Lysosomal Storage Diseases by Providing Positive Feedback Regulation of Lysosomal Ca2+ Release. Developmental Cell, 33(4), 427–441. 10.1016/j.devcel.2015.04.010

Charlton, F. W., Hover, S., Fuller, J., Hewson, R., Fontana, J., Barr, J. N., & Mankouri, J. (2019). Cellular cholesterol abundance regulates potassium accumulation within endosomes and is an important determinant in bunyavirus entry. Journal of Biological Chemistry, 294(18), 7335–7347. 10.1074/jbc.RA119.007618

Cheung, Y. Y., Yu, H., Xu, K., Zou, B., Wu, M., McManus, O. B., Li, M., Lindsley, C. W., & Hopkins, C. R. (2012). Discovery of a Series of 2-phenyl-N-(2-(pyrrolidin-1-yl)phenyl)acetamides as Novel Molecular Switches that Modulate Modes of Kv7.2 (KCNQ2) Channel Pharmacology: Identification of (S)-2-phenyl-N-(2-(pyrrolidin-1-yl)phenyl)butanamide (ML252) as a Potent, B…. Journal of Medicinal Chemistry, 55(15), 6975. 10.1021/JM300700V

Choe, H., Zhou, H., Palmer, L. G., & Sackin, H. (1997). A conserved cytoplasmic region of ROMK modulates pH sensitivity, conductance, and gating. Https://Doi.Org/10.1152/Ajprenal.1997.273.4.F516, 273(4 42–4). 10.1152/AJPRENAL.1997.273.4.F516

Dahal, G. R., Rawson, J., Gassaway, B., Kwok, B., Tong, Y., Ptácek, L. J., & Bates, E. (2012). An inwardly rectifying K+ channel is required for patterning. Development (Cambridge), 139(19), 3653–3664. 10.1242/DEV.078592/-/DC1

Dong, X. P., Shen, D., Wang, X., Dawson, T., Li, X., Zhang, Q., Cheng, X., Zhang, Y., Weisman, L. S., Delling, M., & Xu, H. (2010). PI(3,5)P2 Controls Membrane Traffic by Direct Activation of Mucolipin Ca2+ Release Channels in the Endolysosome. Nature Communications, 1(4), 38. 10.1038/NCOMMS1037

Doyle, D. A., Cabral, J. M., Pfuetzner, R. A., Kuo, A., Gulbis, J. M., Cohen, S. L., Chait, B. T., & MacKinnon, R. (1998). The structure of the potassium channel: Molecular basis of K+ conduction and selectivity. Science, 280(5360), 69–77. 10.1126/SCIENCE.280.5360.69/ASSET/8B1C6173-341A-4492-9124-7C65D5D3CB23/ASSETS/GRAPHIC/SE1586417008.JPEG

Heitmann, A., Gusmag, F., Rathjens, M. G., Maurer, M., Frankze, K., Schicht, S., Jansen, S., Schmidt-Chanasit, J., Jung, K., & Becker, S. C. (2021). Mammals preferred: Reassortment of batai and bunyamwera orthobunyavirus occurs in mammalian but not insect cells. Viruses, 13(9), 1702. 10.3390/V13091702/S1

Hellert, J., Aebischer, A., Haouz, A., Guardado-Calvo, P., Reiche, S., Beer, M., & Rey, F. A. (2023). Structure, function, and evolution of the Orthobunyavirus membrane fusion glycoprotein. Cell Reports, 42(3), 112142. 10.1016/J.CELREP.2023.112142

Hibino, H., Inanobe, A., Furutani, K., Murakami, S., Findlay, I., & Kurachi, Y. (2010). Inwardly Rectifying Potassium Channels: Their Structure, Function, and Physiological Roles. Https://Doi.Org/10.1152/Physrev.00021.2009, 90(1), 291–366. 10.1152/PHYSREV.00021.2009

Hover, S., Charlton, F. W., Hellert, J., Swanson, J. J., Mankouri, J., Barr, J. N., & Fontana, J. (2023). Organisation of the orthobunyavirus tripodal spike and the structural changes induced by low pH and K+ during entry. Nature Communications, 14(1), 1–16. 10.1038/S41467-023-41205-W;TECHMETA

Hover, S., Foster, B., Fontana, J., Kohl, A., Goldstein, S. A. N., Barr, J. N., & Mankouri, J. (2018). Bunyavirus requirement for endosomal K+ reveals new roles of cellular ion channels during infection. PLOS Pathogens, 14(1), e1006845. 10.1371/journal.ppat.1006845

Hover, S., King, B., Hall, B., Loundras, E.-A., Taqi, H., Daly, J., Dallas, M., Peers, C., Schnettler, E., McKimmie, C., Kohl, A., Barr, J. N., & Mankouri, J. (2016). Modulation of Potassium Channels Inhibits Bunyavirus Infection. The Journal of Biological Chemistry, 291(7), 3411–3422. 10.1074/jbc.M115.692673

Ishii, T. M., Silvia, C., Hirschberg, B., Bond, C. T., Adelman, J. P., & Maylie, J. (1997). A human intermediate conductance calcium-activated potassium channel. Proceedings of the National Academy of Sciences of the United States of America, 94(21), 11651. 10.1073/PNAS.94.21.11651

Kendall, R. L., & Holian, A. (2024). Lysosomal BK channels facilitate silica-induced inflammation in macrophages. Inhalation Toxicology, 36(1), 31. 10.1080/08958378.2024.2305112

Krapivinsky, G., Medina, I., Eng, L., Krapivinsky, L., Yang, Y., & Clapham, D. E. (1998). A novel inward rectifier K+ channel with unique pore properties. Neuron, 20(5), 995–1005. 10.1016/S0896-6273(00)80480-8

Lafrenière, R. G., Cader, M. Z., Poulin, J. F., Andres-Enguix, I., Simoneau, M., Gupta, N., Boisvert, K., Lafrenière, F., McLaughlan, S., Dubé, M. P., Marcinkiewicz, M. M., Ramagopalan, S., Ansorge, O., Brais, B., Sequeiros, J., Pereira-Monteiro, J. M., Griffiths, L. R., Tucker, S. J., Ebers, G., & Rouleau, G. A. (2010). A dominant-negative mutation in the TRESK potassium channel is linked to familial migraine with aura. Nature Medicine 2010 16:10, 16(10), 1157–1160. 10.1038/nm.2216

Langereis, M. A., Rabouw, H. H., Holwerda, M., Visser, L. J., & van Kuppeveld, F. J. M. (2015). Knockout of cGAS and STING Rescues Virus Infection of Plasmid DNA-Transfected Cells. Journal of Virology, 89(21), 11169–11173. 10.1128/JVI.01781-15;CTYPE:STRING:JOURNAL

Lewis, L. M., Bhave, G., Chauder, B. A., Banerjee, S., Lornsen, K. A., Redha, R., Fallen, K., Lindsley, C. W., Weaver, C. D., & Denton, J. S. (2009). High-Throughput Screening Reveals a Small-Molecule Inhibitor of the Renal Outer Medullary Potassium Channel and Kir7.1. Molecular Pharmacology, 76(5), 1094. 10.1124/MOL.109.059840

Lopatin, A. N., Makhina, E. N., & Nichols, C. G. (1994). Potassium channel block by cytoplasmic polyamines as the mechanism of intrinsic rectification. Nature, 372(6504), 366–369. 10.1038/372366A0

Lowen, A. C., Noonan, C., McLees, A., & Elliott, R. M. (2004). Efficient bunyavirus rescue from cloned cDNA. Virology, 330(2), 493–500. 10.1016/J.VIROL.2004.10.009

Matsuda, H., Saigusa, A., & Irisawa, H. (1987). Ohmic conductance through the inwardly rectifying K channel and blocking by internal Mg2+. Nature, 325(7000), 156–159. 10.1038/325156A0

Preisig-Müller, R., nter Schlichthö rl, G., Goerge, T., Heinen, S., Brü ggemann, A., Rajan, S., Derst, C., diger Veh, R. W., & rgen Daut, J. (2002). Heteromerization of Kir2.x potassium channels contributes to the phenotype of Andersen’s syndrome. PNAS, 99(11), 7774–7779. www.pnas.orgcgidoi10.1073pnas.102609499

Punch, E. K., Hover, S., Blest, H. T. W., Fuller, J., Hewson, R., Fontana, J., Mankouri, J., & Barr, J. N. (2018). Potassium is a trigger for conformational change in the fusion spike of an enveloped RNA virus. The Journal of Biological Chemistry, 293(26), 9937. 10.1074/JBC.RA118.002494

Reed, A. P., Bucci, G., Abd-Wahab, F., & Tucker, S. J. (2016). Dominant-Negative Effect of a Missense Variant in the TASK-2 (KCNK5) K+ Channel Associated with Balkan Endemic Nephropathy. PLOS ONE, 11(5), e0156456. 10.1371/JOURNAL.PONE.0156456

Rittenhouse, A. R., Parker, C., Brugnara, C., Morgan, K. G., & Alper, S. L. (1997). Inhibition of maxi-K currents in ferret portal vein smooth muscle cells by the antifungal clotrimazole. The American Journal of Physiology, 273(1 Pt 1). 10.1152/AJPCELL.1997.273.1.C45

Sandler, Z. J., Firpo, M. R., Omoba, O. S., Vu, M. N., Menachery, V. D., & Mounce, B. C. (2020). Novel ionophores active against La Crosse virus identified through rapid antiviral screening. Antimicrobial Agents and Chemotherapy, 64(6). 10.1128/AAC.00086-20/FORMAT/EPUB

Scott, C. C., & Gruenberg, J. (2011). Ion flux and the function of endosomes and lysosomes: PH is just the start: The flux of ions across endosomal membranes influences endosome function not only through regulation of the luminal pH. BioEssays, 33(2), 103–110. 10.1002/BIES.201000108;SUBPAGE:STRING:ABSTRACT;WEBSITE:WEBSITE:PERICLES;JOURNAL:JOURNAL:15211878;WGROUP:STRING:PUBLICATION

Shaw, A. B., Tse, H. N., Byford, O., Plahe, G., Moon-Walker, A., Hover, S. E., Saphire, E. O., Whelan, S. P. J., Mankouri, J., Fontana, J., & Barr, J. N. (2024). Cellular endosomal potassium ion flux regulates arenavirus uncoating during virus entry. MBio, 15(7), e01684–23. 10.1128/MBIO.01684-23

Strøbæk, D., Brown, D. T., Jenkins, D. P., Chen, Y. J., Coleman, N., Ando, Y., Chiu, P., Jørgensen, S., Demnitz, J., Wulff, H., & Christophersen, P. (2012). NS6180, a new KCa3.1 channel inhibitor prevents T-cell activation and inflammation in a rat model of inflammatory bowel disease. British Journal of Pharmacology, 168(2), 432. 10.1111/J.1476-5381.2012.02143.X

Strøbæk, D., Hougaard, C., Johansen, T. H., Sørensen, U. S., Nielsen, E., Nielsen, K. S., Taylor, R. D. T., Pedarzani, P., & Christophersen, P. (2006). Inhibitory gating modulation of small conductance Ca2+-activated K+ channels by the synthetic compound (R)-N-(benzimidazol-2-yl)-1,2, 3,4-tetrahydro-1-naphtylamine (NS8593) reduces after hyperpolarizing current in hippocampal CA1 neurons. Molecular Pharmacology, 70(5), 1771–1782. 10.1124/mol.106.027110

Wang, H. R., Wu, M., Yu, H., Long, S., Stevens, A., Engers, D. W., Sackin, H., Daniels, J. S., Dawson, E. S., Hopkins, C. R., Lindsley, C. W., Li, M., & McManus, O. B. (2011). Selective inhibition of the Kir2 family of inward rectifier potassium channels by a small molecule probe: the discovery, SAR and pharmacological characterization of ML133. ACS Chemical Biology, 6(8), 845. 10.1021/CB200146A

Wang, W., Zhang, X., Gao, Q., Lawas, M., Yu, L., Cheng, X., Gu, M., Sahoo, N., Li, X., Li, P., Ireland, S., Meredith, A., & Xu, H. (2017). A voltage-dependent K+ channel in the lysosome is required for refilling lysosomal Ca2+ stores. The Journal of Cell Biology, 216(6), 1715. 10.1083/JCB.201612123

Windhaber, S., Xin, Q., Uckeley, Z. M., Koch, J., Obr, M., Garnier, C., Luengo-Guyonnot, C., Duboeuf, M., Schur, F. K. M., & Lozach, P.-Y. (2022). The Orthobunyavirus Germiston Enters Host Cells from Late Endosomes. Journal of Virology, 96(5). 10.1128/JVI.02146-21

Wu, S. N., Li, H. F., Jan, C. R., & Shen, A. Y. (1999). Inhibition of Ca2+-activated K+ current by clotrimazole in rat anterior pituitary GH3 cells. Neuropharmacology, 38(7), 979–989. 10.1016/S0028-3908(99)00027-1

Wulff, H., Miller, M. J., Hänsel, W., Grissmer, S., Cahalan, M. D., & Chandy, K. G. (2000). Design of a potent and selective inhibitor of the intermediate-conductance Ca2+-activated K+ channel, IKCa1: A potential immunosuppressant. Proceedings of the National Academy of Sciences of the United States of America, 97(14), 8151. 10.1073/PNAS.97.14.8151

Zerr, P., Adelman, J. P., & Maylie, J. (1998). Episodic Ataxia Mutations in Kv1.1 Alter Potassium Channel Function by Dominant Negative Effects or Haploinsufficiency. Journal of Neuroscience, 18(8), 2842–2848. 10.1523/JNEUROSCI.18-08-02842.1998

Zhou, Y., Ä Morais-Cabral, J. H., Kaufman, A., & MacKinnon, R. (2001). Chemistry of ion coordination and hydration revealed by a K + channel±Fab complex at 2.0 A Ê resolution. NATURE, 414. www.nature.com43

