## Supplementary figures and images for "Host cell potassium ion channels KCNJ2 (K_IR_2.1), KCNJ13 (K_IR_7.1) and KCNMA1 (BK_Ca_) mediate escape of Bunyamwera virus from late endosomal compartments"

### Supplemental Figure 1

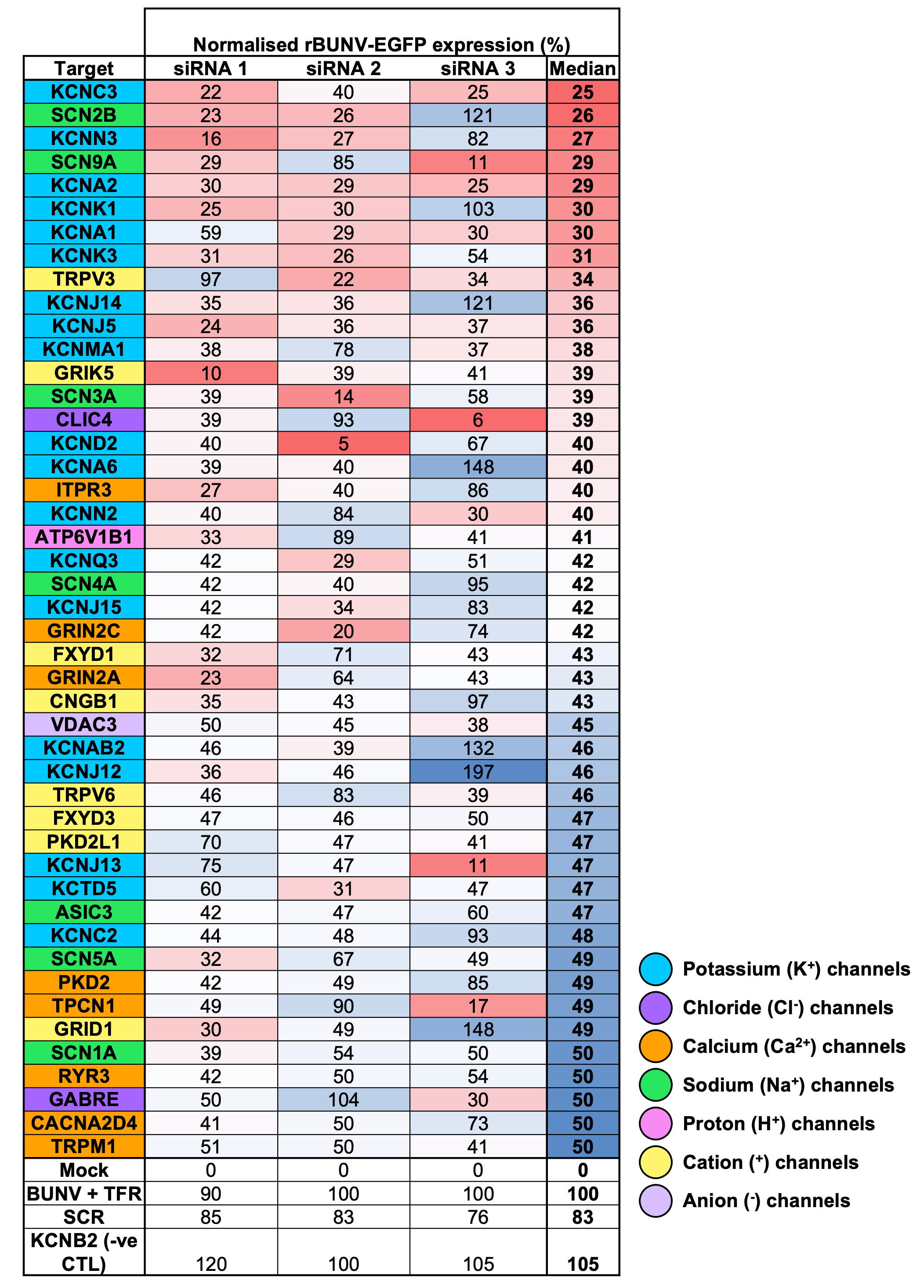

### Supplemental Figure 2

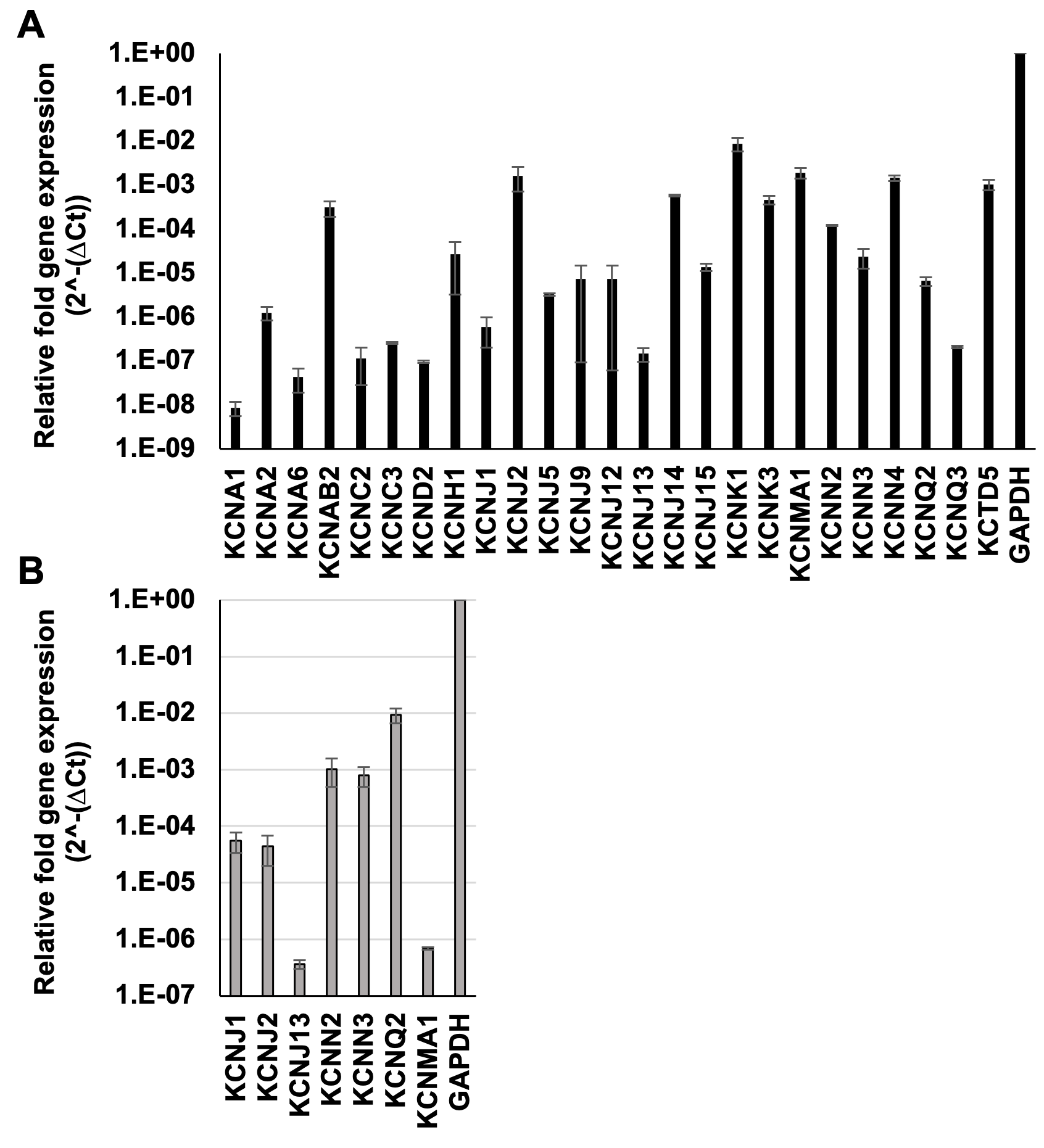

### Supplemental Figure 3

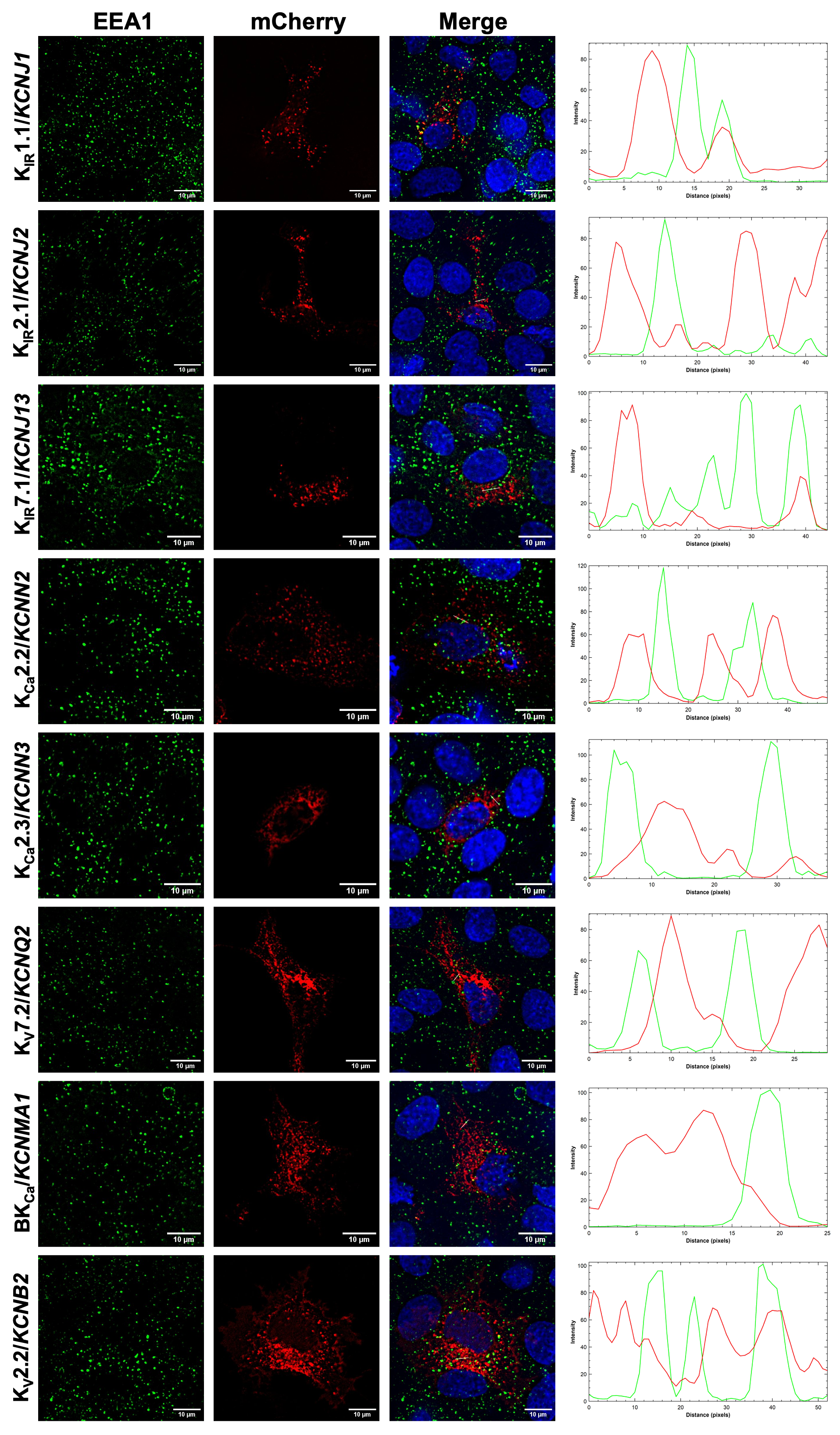

### Supplemental Figure 4

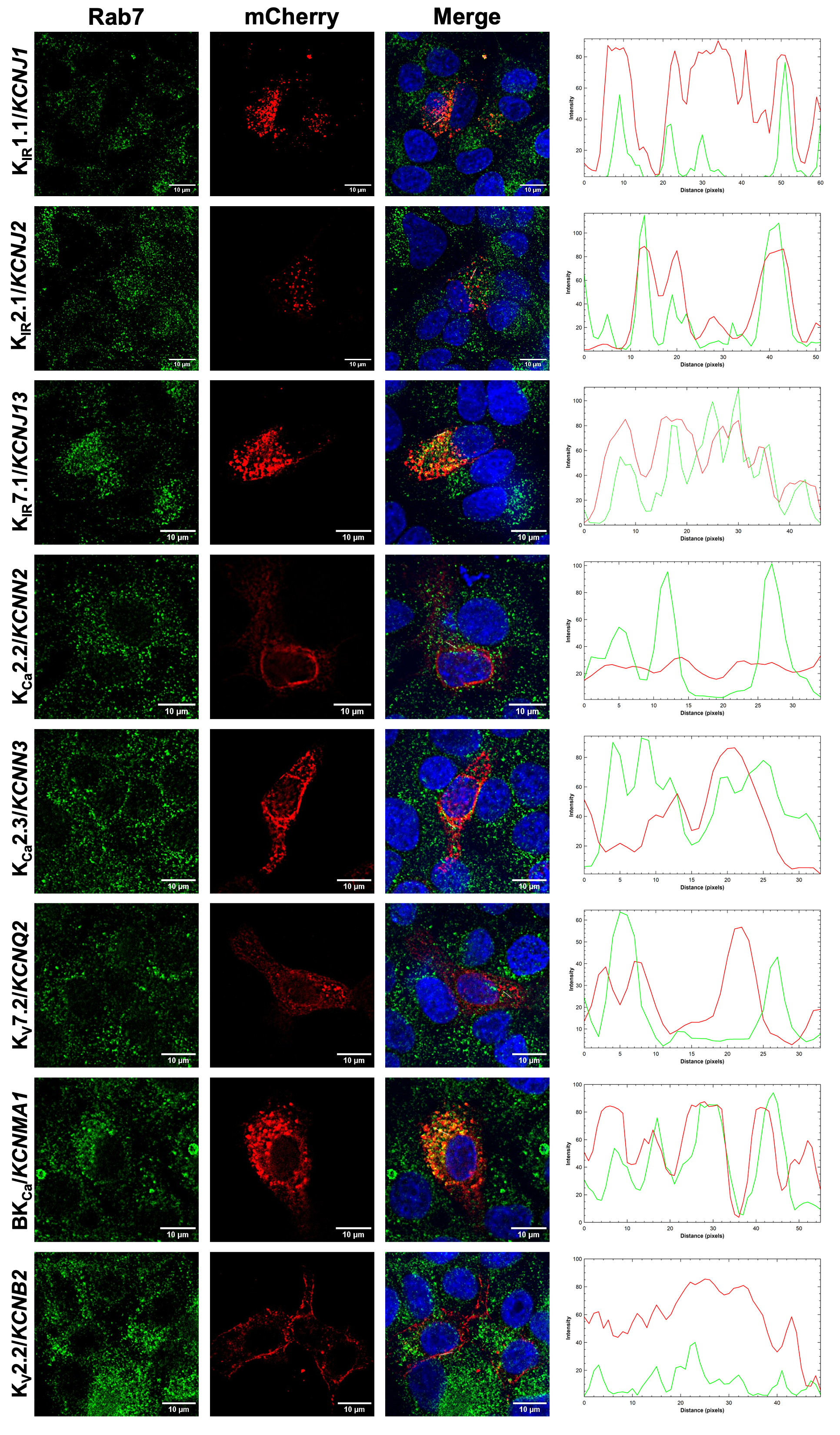
